# FADVI: disentangled representation learning for robust integration of single-cell and spatial omics data

**DOI:** 10.1101/2025.11.03.683998

**Authors:** Wendao Liu, Gang Qu, Lukas M. Simon, Fabian J. Theis, Zhongming Zhao

## Abstract

Integrating single-cell and spatial omics data remains challenging due to strong batch effects across experiments and platforms. Existing methods focus on minimizing these effects but cannot disentangle technical variation from true biological signals. Here, we present FADVI, a variational autoencoder framework partitioning the latent space into batch-specific, label-related, and residual subspaces. By combining supervised classification, adversarial training, and cross-covariance penalty, FADVI disentangles batch from biological representations, preserving biological variation while correcting batch effects. Benchmarking across scRNA-seq, scATAC-seq, and high-resolution spatial transcriptomics datasets, FADVI consistently outperforms state-of-the-art integration methods. FADVI also enables feature attribution for revealing genes associated with cell type identity and batch variation. Together, these results demonstrate that FADVI provides robust, interpretable integration for large-scale single-cell and spatial omics data, offering a powerful framework for downstream analysis and discovery.

## Introduction

With the rapid advancement and widespread adoption of single-cell sequencing and spatial transcriptomics (ST), the scale and complexity of datasets have grown dramatically. Batch effects, caused by unwanted technical variation, arise from sources such as experimental replicates or profiling technologies. These effects present a major challenge for processing atlas-scale datasets and integrating high-resolution ST with single-cell RNA sequencing (scRNA-seq) data. Numerous computational methods have been developed for single-cell batch correction, and comprehensive benchmarking studies have identified top-performing approaches across diverse datasets^1,2^. Among data integration methods, scANVI^3^, a semi-supervised variational autoencoder (VAE)-based method incorporating cell type labels, has consistently achieved state-of-the-art performance, outperforming unsupervised methods.

Spatial transcriptomics has rapidly advanced through parallel developments in both sequencing-based and imaging-based technologies, enabling spatially resolved gene expression profiling in intact tissues. Sequencing-based platforms initially relied on array-based capture with large, multi-cellular spots, but have progressively achieved higher resolution, advancing from technologies such as 10x Genomics Visium to single-cell and subcellular-scale methods like Visium HD^4,5^. In parallel, imaging-based approaches such as Xenium and CosMx^6,7^, which directly visualize transcripts in situ, have matured into highly multiplexed platforms capable of resolving gene expression at single-cell or subcellular resolution. Together, these complementary technological trajectories have greatly expanded the scope of spatially resolved transcriptomic analysis.

Despite these advances, spatial transcriptomics data present distinct analytical challenges. Sequencing-based platforms often exhibit stronger dropout due to limited RNA capture efficiency, while imaging-based platforms typically measure a smaller predefined gene panel. Compared with scRNA-seq, these constraints complicate downstream analyses such as cell type annotation and spatial pattern detection. Integrating spatial transcriptomics data across platforms, or jointly with scRNA-seq, can mitigate these limitations and reveal platform-invariant biological insights, but remains challenging due to differences in resolution, gene coverage, and platform-specific technical effects.

## Results

Unlike existing methods that primarily aim to minimize batch effects, our goal was to disentangle batch variation from biological signals, as disentanglement may improve single-cell data integration, annotation, generation, and counterfactual prediction^8–10^. To this end, we developed FActor Disentanglement Variational Inference (FADVI), a variational autoencoder (VAE)-based framework that partitions the latent space into three distinct subspaces: batch-specific, label-related, and residual variation (Figure 1A).

**Figure 1.**
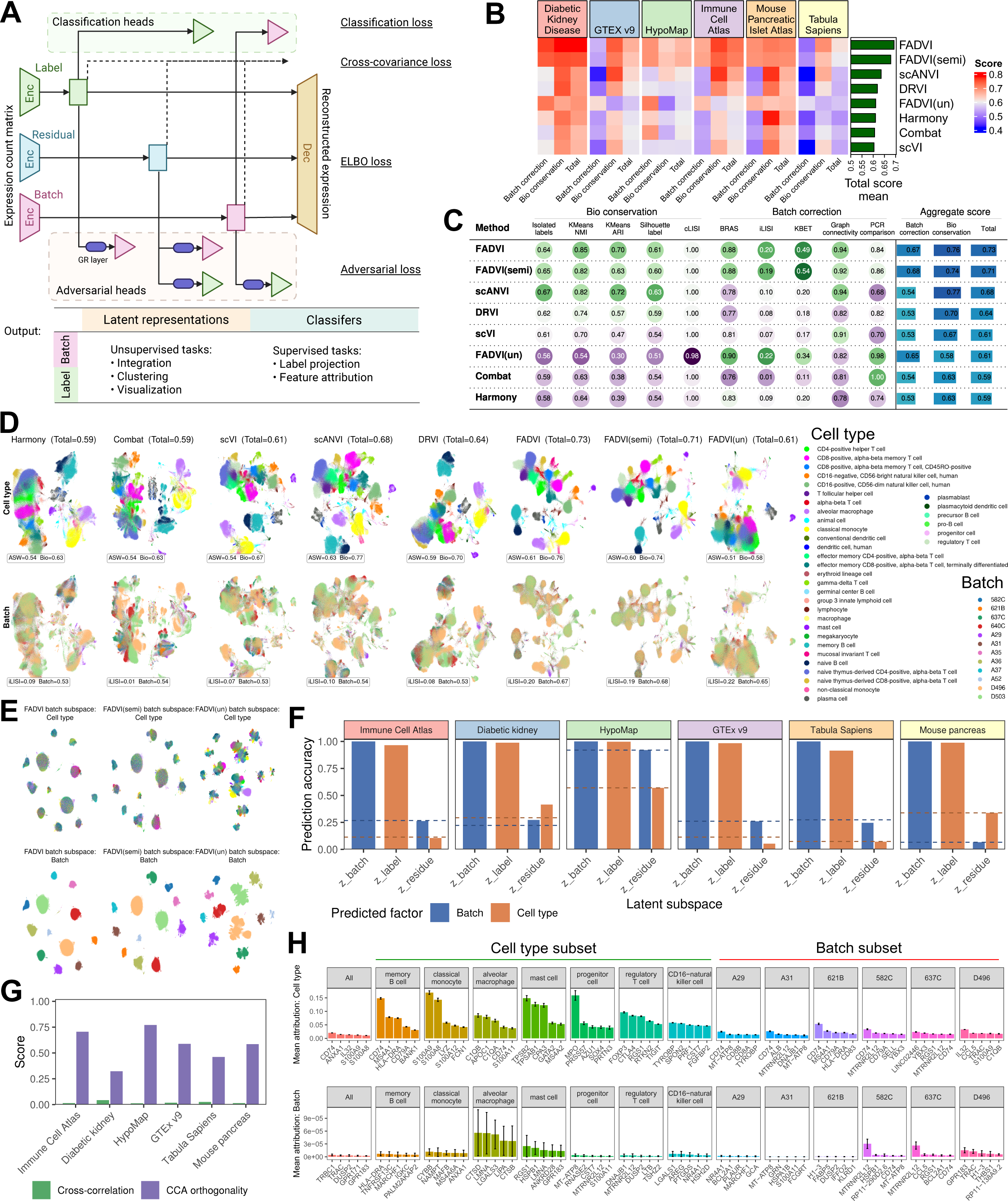
FADVI disentangles batch and biological variation for atlas-scale scRNA-seq data integration. (A) Schematic of FADVI architecture. (B) FADVI shows higher integration scores with competing methods across six scRNA-seq datasets. FADVI(semi): semi-supervised training with masked cell type labels in half batches. FADVI(un): unsupervised training without cell type labels. (C) Individual and aggregated metrics for evaluating integration methods in Immune Cell Atlas. (D) UMAP visualization of cell type labels and batches across integration methods using Immune Cell Atlas dataset, annotated with average silhouette width (ASW), integration local inverse Simpson’s index (iLISI), and aggregate scores. (E) UMAP plots of cell type labels and batches using FADVI representations in batch subspace. (F) Accuracy of predicting batch and cell type from each cell representation in latent subspaces by cross-validated classification. Dashed lines indicate the majority-class baseline for each factor. (G) Mean absolute cross-subspace correlation and canonical-correlation orthogonality. (H) Prioritized genes associated with cell type labels or batch effect in the whole dataset, individual cell types, and batches. High attribution variance suggested substantial expression variability across batches. Error bars represent standard error.

The encoder maps input into three latent factors, which are then combined by the decoder to reconstruct the original data. To ensure that each factor captures only the intended information, we introduce supervised classification heads, adversarial networks with gradient reversal layers, and a cross-covariance penalty. The supervised heads align the batch and label factors with input observations, while the adversarial heads prevent leakage of unintended information across factors. The cross-covariance penalty further enforces statistical independence among the latent subspaces. This design enables FADVI to generate disentangled representations that support integration, clustering, visualization, and label projection, while enhancing interpretability and robustness. Similar to scANVI, FADVI follows a semi-supervised paradigm, capable of integrating both labeled and unlabeled data with shared cell type composition. Unless otherwise stated, all integration results and Uniform Manifold Approximation and Projection (UMAP) visualizations are derived from the label-related subspace, which is the representation we recommend for integration, clustering and visualization; the batch subspace is shown separately where indicated.

We benchmarked FADVI with other integration methods on three tasks: atlas-scale scRNA-seq integration, single-cell ATAC sequencing (scATAC-seq) integration, and high-resolution ST with paired scRNA-seq integration. The top-performing methods from two consecutive benchmarking studies were selected for benchmarking, including semi-supervised scANVI^3^, and unsupervised scVI^11^, Combat^12^, Harmony^13^, and LIGER^14^. We also included another single-cell data disentanglement method DRVI^15^.

For scRNA-seq integration, we evaluated FADVI on six datasets from a recent benchmarking study^1^: diabetic kidney disease dataset^16^, GTEX v9^17^, HypoMap^18^, Immune Cell Atlas^19^, Mouse Pancreatic Islet Atlas^20^, and Tabula Sapiens^21^. Across all datasets, both fully-supervised FADVI (with complete cell type labels) and semi-supervised FADVI (with partial labels) consistently outperformed competing methods, achieving substantially higher batch correction scores while maintaining comparable levels of biological variation conservation (Figure 1B). Unsupervised FADVI had similar performance as other unsupervised methods. For instance, in Immune Cell Atlas with 329,762 cells from 12 batches, FADVI and semi-supervised FADVI improved the aggregated batch correction score by approximately 26% compared to other methods (Figure 1C). Latent space visualization using UMAP further highlighted FADVI’s advantage, displaying better separation of cell clusters alongside improved mixing of batches (Figure 1D). In addition, when projecting into the batch-specific subspace, FADVI produced representations where batch-specific clusters were clearly distinguishable while cell types were intermixed within each batch (Figure 1E), the mirror image of the integration embedding and a direct confirmation that technical and biological variation were successfully disentangled. Similar patterns were observed across all scRNA-seq datasets (Figure S1-S5). Moreover, the classification head in semi-supervised FADVI accurately predicted cell types for unlabeled cells, demonstrating its effectiveness in capturing biological variation even with incomplete label information (Figure S6). To test the factorization directly rather than by proxy, we quantified how well each factor could be recovered from each subspace by cross-validated classification (Figure 1F, Table S1). Batch identity was predicted from batch representation (**z**_b_) with accuracy above 0.999 in all six datasets, and cell type from cell representation (**z**_l_) with accuracy between 0.915 and 0.998, whereas the reverse assignments and predictions from the residual representation (**z**_r_) remained close to the majority-class baseline. One exception is informative: in HypoMap, 92.0% of cells derive from a single batch and 57.0% are neurons, so the apparently high accuracy of predicting batch from **z**_r_ simply matches the majority-class rate and reflects the dataset’s extreme composition rather than information leakage. Mean absolute cross-correlation between subspaces was 0.012–0.043, and canonical-correlation analysis indicated medium-to-high orthogonality that varied across datasets (0.32– 0.77, Figure 1G). Collectively, these results demonstrate that batch and biological information concentrate in their intended subspaces, and that **z**_r_ does not act as a sink for either signal. Because FADVI is semi-supervised, we further questioned how little supervision it needs. Reducing the labeled fraction from 100% to 1% retained 88–99% of fully-supervised integration quality across all nine datasets, and FADVI exceeded scANVI at every labeled fraction in most of the nine datasets (Figure S7, Table S2). Performance remained stable when the labeled and unlabeled subsets had markedly different cell type compositions, or when 5–20% of labels were corrupted. This suggested the robust utility of FADVI when annotations are sparse, imbalanced or imperfect (Figure S7).

Leveraging FADVI’s ability to disentangle batch and biological variation, we applied Shapley Additive Explanations (SHAP) to identify genes associated with cell type labels and batch effects^22^. In Immune Cell Atlas, known marker genes were consistently prioritized for label prediction, while distinct sets of genes were associated with batch effects in different cell types, such as *CTSD* and *LMNA* in alveolar macrophage, and *RGS1* and *HSPB1* in mast cells (Figure 1H). We also detected batch-specific signatures, such as *TRBC1* and *TRAC* for overall batch effects, *MTRNR2L12* in batches 582C/637C and *GPR183* in D496, which showed large variation across different batches (Figure S8). Because feature attribution is interpretable only when a subspace captures the intended signal, we repeated the attribution analysis separately for each of the three subspaces using integrated gradients on the latent norm (Figure S9, Table S3). Genes attributed to **z**_b_ and **z**_l_ were largely distinct, while attributions to **z**_r_ had little overlap with either, consistent with the predictability results above.

We evaluated the contribution of individual loss components in FADVI under supervised and semi-supervised settings (Figure S10, Table S4). Removing label classification losses caused the largest decline, underscoring the essential role of supervised signal for representation learning. Adversarial terms were also beneficial, with complete removal leading to strong deterioration in batch correction. Among individual components, residual adversarial losses had minimal effect, while cross batch–label terms provided moderate benefits. The Cross-covariance loss, although not critical for predictive accuracy, contributed to more effective batch effect correction. Moreover, batch-level masking was generally more robust than stratified label retention for semi-supervised models. Because each ablation sets a single loss weight to zero, we additionally conducted a one-at-a-time sensitivity analysis over the full range of weights (Figure S11, Table S5). Across six scRNA-seq datasets, the total integration score varied by only 0.015–0.041 over KL divergence weight (β) in [0.1, 5], batch classifier loss weight (λ_b_) or label classifier loss weight (λ_l_) in [10, 200], and gradient reversal weight (α) or cross-covariance weight (γ) in [0, 10]. This limited variation suggests that the default settings are near-optimal without the need for dataset-specific tuning. The observed trends are smooth and interpretable: increasing β shifts the trade-off from biological conservation toward batch correction, whereas increasing λ_l_ enhances biological conservation, and the adversarial and cross-covariance weights have the smallest overall effect.

For scATAC-seq integration, we evaluated FADVI using mouse brain datasets from a previous benchmarking study^2^, including the full dataset (Large ATAC) and a downsampled version (Small ATAC). Each dataset was represented in three feature spaces: peaks, windows and gene activity. Peaks are the most commonly used features for ATAC-seq, representing small open chromatin regions. Windows were equal-length non-overlapping regions across the genome. Gene activity was calculated using peaks in the gene body and promoter region. Both FADVI and semi-supervised FADVI outperformed competing methods, including LIGER, the top-ranked method in the original benchmarking (Figure 2A)^2^. In particular, even though scANVI is a supervised method, its performance was lower than unsupervised LIGER and semi-supervised FADVI, suggesting that method design could be more important than supervision for the scATAC-seq data integration task. UMAP visualization further confirmed the disentangled variations across three feature spaces (Figure S12-S17).

**Figure 2.**
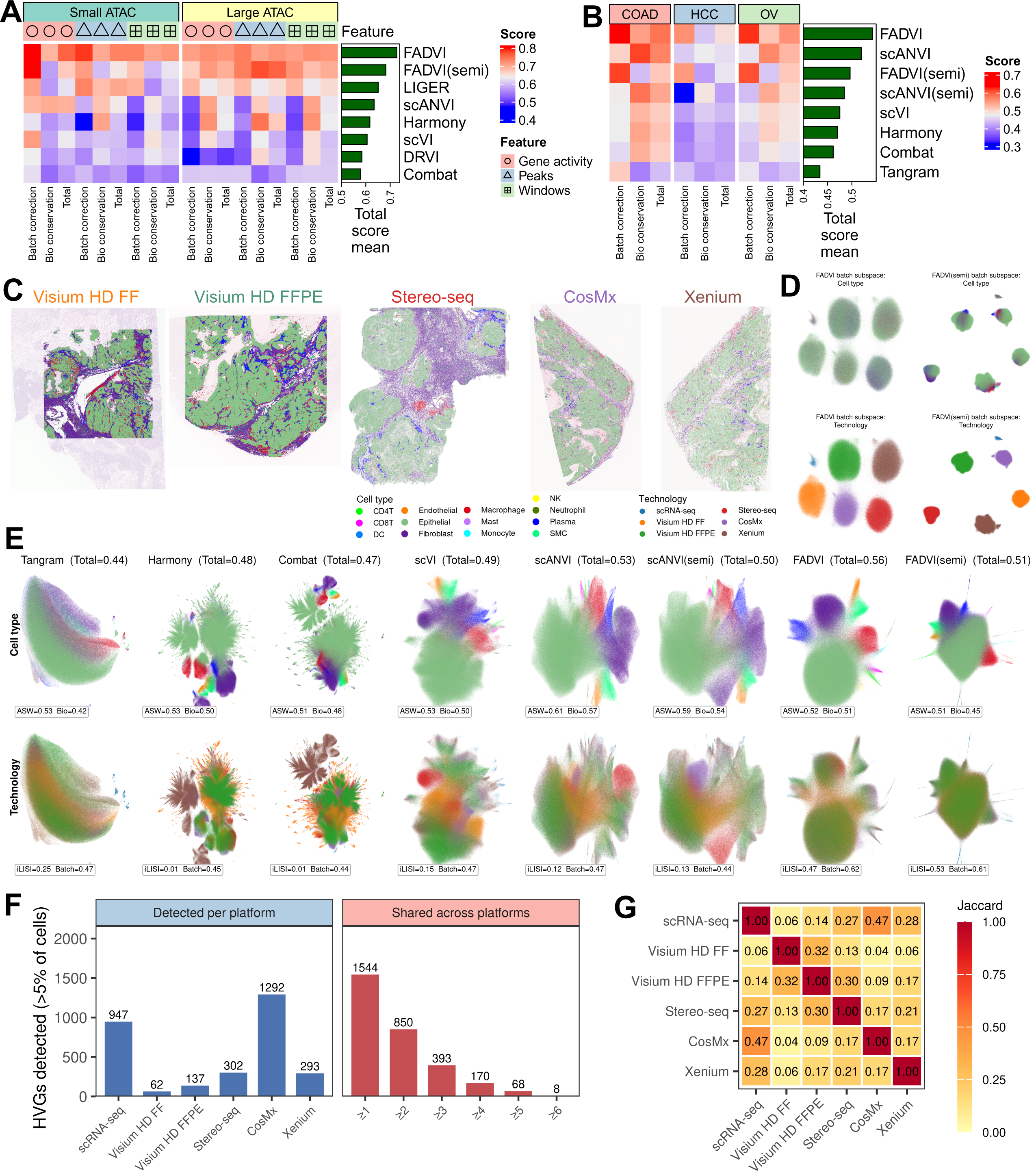
FADVI enables robust integration of scATAC-seq data, and high-resolution ST across diverse platforms with scRNA-seq data. (A) FADVI shows higher integration scores with competing methods across two scATAC-seq datasets. (B) FADVI shows higher integration scores for integrating ST with scRNA-seq data across three cancer types. FADVI(semi) and scANVI(semi): semi-supervised training with masked cell type labels in all ST data. (C) Cell type labels of OV ST data overlaid on hematoxylin and eosin staining image. FF: fresh frozen; FFPE: Formalin-Fixed Paraffin-Embedded. (D) UMAP plots of cell type labels and batches in OV ST data using FADVI representations in batch subspace. (E) UMAP visualization of cell type labels and batches across integration methods, annotated with ASW, iLISI, and aggregate scores. (F) The number of 2,000 highly variable genes (HVGs) detected (in > 5% of cells) on each platform in the OV dataset (left), and the number shared by at least k platforms (right). On the right panel, it shows the feature space common to all six platforms is almost empty. (G) Pairwise Jaccard overlap between platforms’ detected-gene sets in the OV dataset.

The most challenging task was integrating high-resolution ST across diverse platforms with paired scRNA-seq data. We analyzed three datasets profiling colon adenocarcinoma (COAD), hepatocellular carcinoma (HCC), and ovarian cancer (OV), generated using five ST platforms alongside scRNA-seq^23^. Among them, CosMx, Stereo-seq, and Xenium provide single-cell resolution, whereas Visium HD captures 8 μm spots. In the semi-supervised setting, all ST cells or spots were treated as unlabeled, representing >99% of the data. For comparison, we also benchmarked Tangram^24^, a method designed for aligning ST with scRNA-seq data. Semi-supervised methods (FADVI and scANVI) outperformed unsupervised methods, while fully-supervised methods with complete cell type labels achieved the best performance (Figure 2B). Notably, FADVI surpassed scANVI in both semi- and fully-supervised contexts. Considering technologies as batches (Figure 2C), FADVI effectively captured technology-associated variation (Figure 2D). UMAP visualization of the integrated representations indicated poor batch mixing in all methods except FADVI (Figure 2E). However, semi-supervised FADVI also revealed limited integration between scRNA-seq and ST data, suggesting that scRNA-seq labels alone were insufficient to fully guide integration (Figure 2E, Figure S18-S19). To understand why cross-platform ST integration is inherently challenging, we quantified the extent of platforms divergence in the feature space used for integration (Table S6). Among the 2,000 highly variable genes used for integration in the OV dataset, the number detected in more than 5% of cells varied approximately 20-fold across platforms (62 for Visium HD FF to 1,292 for CosMx), and the number detected on all six platforms was only 8 (0.4%) (Figure 2F). Pairwise Jaccard overlap among platforms’ detected-gene sets was correspondingly low (0.04–0.47), with the lowest overlap between imaging- and sequencing-based assays (Figure 2G). The same pattern held in COAD (32 of 2,000 shared, 1.6%) and HCC (6 of 2,000, 0.3%) (Figure S20). Because the shared feature space is nearly empty, methods that assume a common feature distribution have little to align, explaining the platform-specific failures we observed: when quantifying per-platform mixing across methods, Xenium was the most common point of failure, and FADVI was the only method that never failed on any platform-dataset combination (Figure S21, Table S7).

We further compared FADVI against two spatially-aware integration methods that explicitly use coordinates, GraphST and STAligner^25,26^. Because neither method could be run on the full datasets within the available memory, we compared all four methods using contiguous center spatial crops of approximately 12,500 cells per platform. This approach preserves local adjacency and thus, the information on which these methods rely. FADVI achieved a higher total score than both spatially-aware methods on all three cancer types (FADVI mean: 0.617, STAligner: 0.370, and GraphST: 0.317), with the advantage driven primarily by batch correction while biological conservation remained comparable (Figure S22, Table S8). This indicates that, for the specific task of aligning high-resolution ST with scRNA-seq across platforms, disentangling technical from biological variation in expression space is more effective than incorporating spatial coordinates. Subspace predictability in the three ST datasets mirrored the scRNA-seq results: batch identity was recovered from **z**_b_ with accuracy exceeding 0.999, while cell type was recovered from **z**_l_ with an accuracy of 0.929–0.967, cross-subspace correlation ranged from 0.006 to 0.009, and again **z**_r_ showed no evidence of functioning as a signal sink (Table S1).

## Discussion

We introduce FADVI, a generative neural network that leverages disentangled representation learning to integrate single-cell and spatial transcriptomics data. Across diverse scRNA-seq, scATAC-seq, and spatial transcriptomics datasets, FADVI consistently achieved robust integration and outperformed state-of-the-art methods. Although both FADVI and scANVI are based on a variational autoencoder architecture and a semi-supervised paradigm, FADVI more effectively removes batch effects by explicitly modeling batch-specific variation in a dedicated latent subspace. By disentangling technical and biological signals, FADVI not only enhances integration performance but also enables feature prioritization associated with cell type identity and batch effects. FADVI’s conceptual advance lies not in any established single component, but in its explicit three-way latent-space factorization and its redundant enforcement through supervised heads, adversarial heads, and a non-adversarial decorrelation penalty. scVI and scANVI learn a single latent space in which batch effects are marginalized rather than represented. Thus, technical variation cannot be inspected or attributed. By assigning batch variation to a dedicated subspace, FADVI improves batch correction while preserving the removed variation for downstream analysis, enabling the batch-specific feature attribution reported above.

FADVI has several limitations. First, it is designed primarily as a semi-supervised method. While the availability of partial labels substantially improves its performance, FADVI does not outperform competing approaches in fully unsupervised settings. Second, similar to many existing integration methods, FADVI does not explicitly resolve confounding between biological and technical factors. When biological conditions of interest—such as perturbations or disease states—are correlated with batch effects, relevant biological signals may be absorbed into the batch-specific subspace. Consequently, careful experimental design and data collection remain essential to minimize confounding and ensure accurate biological interpretation. Third, for spatial datasets, FADVI operates on expression profiles and does not use spatial coordinates as input. In benchmarking, FADVI outperformed two spatial coordinate-based methods (GraphST and STAligner) on the cross-platform integration with its strength in batch correction. Accordingly, we consider the cross-platform integration problem addressed in this work as one primarily driven by technical, rather than spatial, variation. However, tasks defined intrinsically in space, such as spatial domain identification, will require models that use coordinates directly. Finally, FADVI is more computationally demanding than competing VAE-based methods. It requires roughly 3.5 times the per-epoch training time of scVI and 1.4 times that of scANVI. This reflects the cost of its four auxiliary classification heads and the cross-covariance penalty. Overall, the absolute cost remains modest: integrating 300,000 cells takes under one hour on a single GPU, runtime scales linearly with cell number, and GPU memory usage is independent of dataset size due to mini-batched training (Table S9).

We anticipate that FADVI will provide a powerful framework for integrating large-scale single-cell and spatial omics datasets, particularly when batch effects are strong. Future work may extend disentangled representation learning to transformer-based foundation models for single-cell data. Considering that most existing single-cell foundation models do not yet achieve better integration performance when compared with negative controls^1^, explicitly disentangling batch-related and biological variation during self-supervised pretraining or downstream fine-tuning would substantially enhance the quality of learned cell embeddings and improve cross-dataset integration.

## Resource availability

### Lead contact

Further information and requests for resources should be directed to and will be fulfilled by the lead contact, Zhongming Zhao.

### Materials availability

This study did not generate new materials.

### Data and code availability

- This paper analyzes existing, publicly available data. Public datasets were available at: https://openproblems.bio/datasets (scRNA-seq), https://doi.org/10.6084/m9.figshare.12420968 (scATAC-seq), https://spatch.pku-genomics.org/#/download (ST with paired scRNA-seq).
- FADVI is publicly available at https://github.com/liuwd15/fadvi, and the documentation is available at https://fadvi.readthedocs.io.

## Acknowledgements

This work was partially supported by National Institutes of Health grants U01AG079847 and R01LM012806. We thank the technical support of the Cancer Genomics Core supported by the Cancer Prevention & Research Institute of Texas (CPRIT) grant RP240610. W.L. was supported by John and Rebekah Harper Fellowship in Biomedical Sciences. W.L. and G.Q. were supported by CPRIT Fellowship in the Biomedical Informatics, Genomics and Translational Cancer Research Training Program (BIGTCR, CPRIT RP210045). The funders had no role in the study design, data collection and analysis, or decision to publish or prepare the manuscript.

## Author contributions

W.L. conceived the project, developed FADVI method, and performed most analyses. G.Q. performed ablation analysis. L.M.S., F.J.T., and Z.Z. contributed to method evaluation. Z.Z. supervised the project. All authors wrote and reviewed the manuscript.

## Declaration of interests

W.L. joined Bristol Myers Squibb during the peer review process. Bristol Myers Squibb had no role in the study design, data collection and analysis, or the decision to publish. F.J.T. consults for Immunai Inc., CytoReason Ltd, Cellarity, BioTuring Inc., and Genbio.AI Inc., and has an ownership interest in Dermagnostix GmbH and Cellarity. The remaining authors declare no competing interests.

## Declaration of generative AI and AI-assisted technologies in the writing process

During the preparation of this work, the authors used Claude Code in order to improve manuscript writing. After using this tool, the authors reviewed and edited the content as needed and take full responsibility for the content of the published article.

## Supplementary information

Document S1. Supplemental Figures S1-S22.

Document S2. Supplemental Tables S1-S9.

## STAR methods

### Key resources table

| REAGENT or | SOURCE | IDENTIFIER |
| --- | --- | --- |

| RESOURCE |  |  |
| --- | --- | --- |
| **Deposited data** |  |  |
| Diabetic kidney disease scRNA-seq data | Wilson et al.<sup>16</sup> | https://openproblems.bio/datasets/cellxgene_census/dkd |
| GTEx v9 scRNA-seq data | Eraslan et al.<sup>17</sup> | https://openproblems.bio/datasets/cellxgene_census/gtex_v9 |
| HypoMap scRNA-seq data | Steuernagel et al.<sup>18</sup> | https://openproblems.bio/datasets/cellxgene_census/hypomap |
| Immune Cell Atlas scRNA-seq data | Dominguez Conde et al.<sup>19</sup> | https://openproblems.bio/datasets/cellxgene_census/immune_cell_atlas |
| Mouse Pancreatic Islet Atlas scRNA-seq data | Hrovatin et al.<sup>20</sup> | https://openproblems.bio/datasets/cellxgene_census/mouse_pancreas_atlas |
| Tabula Sapiens scRNA-seq data | Jones et al.<sup>21</sup> | https://openproblems.bio/datasets/cellxgene_census/tabula_sapiens |
| Mouse brain scATAC-seq data | Luecken et al.<sup>2</sup> | https://doi.org/10.6084/m9.figshare.12420968 |
| Human cancer ST data | Ren et al.<sup>23</sup> | https://spatch.pku-genomics.org/#!/download |
| **Software and algorithms** |  |  |
| scib (v1.1.7) | Luecken et al.<sup>2</sup> | https://github.com/theislab/scib |
| scib-metrics (v0.5.6) | Luecken et al.<sup>2</sup> | https://github.com/yoseflab/scib-metrics |
| scvi-tools (v1.4.3) | Lopez et al.<sup>11</sup> | https://github.com/scverse/scvi-tools |
| Harmony(v0.0.10) | Korsunsky et | https://github.com/slowkow/harmonypy |
|  | al.<sup>13</sup> |  |
| DRVI (v0.1.9) | Moinfar et al.<sup>15</sup> | https://github.com/theislab/DRVI |
| LIGER (v2.2.1) | Welch et al.<sup>14</sup> | https://github.com/welch-lab/liger |
| Tangram (v1.0.4) | Biancalani et al.<sup>24</sup> | https://github.com/broadinstitute/Tangram |
| Scanpy (v1.12.1) | Virshup et al.<sup>27</sup> | https://github.com/scverse/scanpy |
| GraphST(v1.1.1) | Long et al.<sup>25</sup> | https://github.com/JinmiaoChenLab/GraphST |
| STAligner(v1.0.0) | Zhou et al.<sup>26</sup> | https://github.com/zhoux85/STAligner |
| FADVI (v0.3.0) | This work | https://github.com/liuwd15/fadvi/ |

## Method Details

### Model architecture

FADVI explicitly separates latent representations into three parts: label-related factors (**z**_l_), <u>b</u>atch-related factors (**z**_b_), and residual factors (**z**_r_). Given an input observation ***x*** ∊ R^D^, the encoder produces Gaussian parameters for each latent subspace:

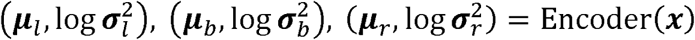

By default, we use the raw read count as input ***x***, which is assumed to follow the zero-inflated negative binomial (ZINB) distribution.

Each latent is then sampled using the reparameterization trick:

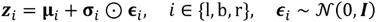

The concatenated latent representation is decoded back into the input space:

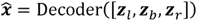

The VAE is trained with the evidence lower bound (ELBO):

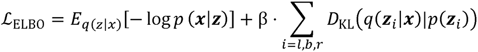

where *p*(**z**_i_)is the standard Gaussian prior, and β controls KL regularization strength. The first term corresponds to the reconstruction loss.

To enforce semantic alignment, fully-connected classification heads are attached with supervised cross-entropy (CE) loss:

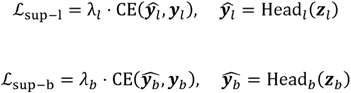

To ensure each subspace is exclusive, fully-connected adversarial heads are trained via gradient reversal (GR) to prevent leakage: Importantly, the gradient reversal layer is the identity in the forward pass and negates and rescales the gradient in the backward pass, so the encoder and the auxiliary classifiers are optimized jointly in a single backward pass under ordinary stochastic gradient descent. Accordingly, FADVI avoids alternating minimax optimization between separately updated networks; the setting in which adversarial training is susceptible to oscillation and mode collapse. The adversarial weights are small and fixed (α = 1 by default, never scheduled) relative to the reconstruction and supervision terms (λ = 50), and the cross-covariance penalty targets the same objective without adversarial dynamics. In this case, disentanglement does not depend on the adversary alone.

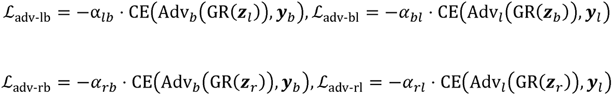

To further decorrelate the latent subspaces, we apply a cross-covariance penalty to minimize statistical dependence between different latent factors:

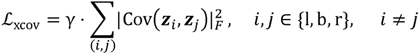

The total loss combines all components:

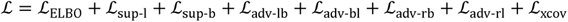

For unlabeled cells, *L*_sup-l_ , *L*_adv-bl_ , *L*_adv-rl_ are set to 0.

### Batch integration benchmarking

We benchmarked FADVI against several integration methods: scANVI^3^, scVI^11^, Combat^12^, Harmony^13^ from Python package scib (v1.1.7)^2^; LIGER from R package rliger (v2.2.1)^14^; DRVI from Python package drvi-py (v0.1.9)^15^; Tangram from Python package Tangram (v1.0.4)^24^. Running scDisco failed on all datasets so it is not included^8^. All methods except DRVI were run with default parameters. For scRNA-seq datasets, DRVI was run using additional parameters: {n_latent=128, max_epochs=400, early_stopping=False, n_epochs_kl_warmup=400}. For scRNA-seq and ST–scRNA-seq integration, the top 2,000 highly variable genes were used as input, while for scATAC-seq integration, all features (peaks, windows, or gene activity) were included. Latent representations generated by each method were evaluated using scib_metrics (v0.5.6) and visualized with UMAP using Scanpy (v1.11.1)^27^.

FADVI and scANVI were tested under supervised, semi-supervised, and unsupervised settings. In the supervised setting, all cell type labels were provided as input, whereas in the unsupervised setting no labels were used. The semi-supervised setting simulated real-world scenarios by masking labels in specific subsets: half batches in scRNA-seq datasets, two smaller batches in scATAC-seq datasets, and all ST data in the paired ST-scRNA-seq datasets. For benchmarking, we used label subspace representations from supervised and semi-supervised FADVI, and the concatenated label and residual subspaces from unsupervised FADVI.

### Subspace disentanglement quantification

To quantify disentanglement directly, we trained a multinomial logistic-regression classifier to predict batch and cell type labels from each latent subspace (**z**_b_, **z**_l_, **z**_r_) and report five-fold cross-validation accuracy, ensuring that the values are not inflated by overfitting. Accuracies are interpreted against the majority-class rate rather than against 1/n_classes, as class frequencies are highly non-uniform in several datasets. Statistical dependence between subspaces was summarized as the mean absolute Pearson correlation over all pairs of dimensions, and as a canonical-correlation-based orthogonality score. We also evaluated whether **z**_r_ acts as a signal sink by comparing its batch and cell type accuracy values predicted from **z**_r_ with the corresponding majority-class baselines, and by comparing its explained variance fraction with those of **z**_b_ and **z**_l_. Feature attributions were computed per subspace with integrated gradients applied to the L2 norm of each subspace, and were residualized against expression rank before comparison, because raw attribution magnitude correlates strongly with expression level.

### Label dependency

To assess dependence on labeled data, cell type labels were retained for 1%, 5%, 10%, 20%, 50% and 100% of cells by stratified sampling within each batch, with all remaining cells treated as unlabeled, and the resulting representations were scored with scib-metrics. Two additional variants were tested: label noise – 5%, 10% or 20% of retained labels were permuted, and imbalanced composition – abundant cell types were labeled at 1% while the remaining types were labeled at 50% so that labeled and unlabeled subsets differ markedly in composition. For the imbalanced experiment, a balanced control using the same total number of labels, drawn proportionally across cell types, was trained for comparison. Semi-supervised scANVI was run under the identical label masks at every condition. Unsupervised scVI was included as a no-label reference.

### Model prediction interpretability

FADVI implements interpretability analysis using GradientShap from Captum to explain model predictions at the feature level^22^. It computes feature attributions by analyzing how each gene contributes to batch or label classification decisions through gradient-based methods. Batch predictions trace gradients through the batch subspace, while label predictions use the label subspace. Cell-level attributions are then transformed into feature-level importance. For each gene, FADVI computes mean attribution, standard deviation, and mean absolute attribution. High mean attribution or mean absolute attribution indicate features strongly associated with cell type labels or batch effect.

### Ablation study

To evaluate the role of individual loss components, we conducted ablation experiments under supervised and semi-supervised settings. The semi-supervised setting was same as the benchmarking setting by masking labels in half batches in scRNA-seq datasets. The network architecture was fixed across all experiments, and ablated models were created by selectively removing specific objectives: cross-covariance loss, label classification, batch classification, all adversarial terms, residual-subspace adversarial losses, or cross batch-label adversarial losses. Integration performance was assessed using biological conservation, batch correction, and total scores.

### Hyperparameter sensitivity and training stability

Hyperparameter sensitivity was evaluated using the semi-supervised FADVI model, with cell type labels masked in half of the batches same as the benchmark settings. Each loss weight varied by one factor at a time from the default configuration while all other weights were held at their defaults, sweeping β over [0.1, 5], λ_b_ and λ_l_ over [10, 200], and α and γ over [0, 10]; a single baseline model was trained once and used as the common reference point for every curve, enabling direct comparison across all sweeps. Because setting a weight to zero is equivalent to removing the corresponding loss term, these sweeps extended the ablation study described above. Training stability under gradient reversal was examined separately by sweeping the reversal strength α over 0, 0.5, 1, 5, 10 and 50 and then recording each loss component per epoch.

### Comparison with spatially-aware integration methods

GraphST (v1.1.1) and STAligner (v1.0.0) were run on the three ST datasets using spatial coordinates as intended by each method: GraphST constructs its neighbor graph and adjacency matrix from obsm[‘spatial’], and STAligner computes a spatial network per slice with a radius cutoff of 50 units before training. Because neither method could process the full datasets within the available memory, we compared all methods using contiguous square crops from the center of each platform’s tissue section, comprising approximately 12,500 cells per platform. Contiguous cropping rather than random subsampling is required here, because random subsampling would destroy the local adjacency structure on which these methods rely. The scRNA-seq reference was excluded from this comparison due to lack of its coordinates. All cell representations were benchmarked using scib-metrics. FADVI used the same hyperparameters as in the main benchmark.

### Computational efficiency

Training time and memory usage were measured for FADVI, scVI and scANVI on 10,000, 50,000, 100,000 and 300,000 cells subsampled from the Immune Cell Atlas, using 2,000 highly variable genes, a batch size of 256, and a single NVIDIA L40S GPU. Per-epoch time was reported as the median over a fixed epoch budget; thus, one-off first-epoch overhead did not distort it. Time to convergence was determined post hoc from the recorded validation-ELBO curve using one deterministic rule applied identically to every method: convergence is the first epoch after which five consecutive epochs lie within 2% of the best value reached in that run. Because integration quality could plateau before the loss does, we additionally snapshotted the cell embedding every 10 epochs over 100 epochs and scored each snapshot with scib-metrics, defining convergence as the first checkpoint within 0.01 of that method’s own best total score. Peak GPU memory was measured using torch.cuda.max_memory_allocated, and peak host memory defined as the maximum resident set size.

## Notes

### Summary of Updates

Major improvements in this revision: We quantified the disentanglement of the three latent subspaces directly, measuring how well batch and cell type can be recovered from each subspace across six scRNA-seq and three ST datasets, and reporting subspace independence by cross-correlation and canonical-correlation analysis (Figure 1F, 1G; Table S1). We tested the residual subspace for signal leakage and expanded feature attribution to each subspace separately, confirming that the residual subspace is not a sink for batch or biological signal (Figures S9; Tables S1, S3). We characterized FADVI's dependence on labels, showing that 1% of labels could retain 88-99% of fully-supervised performance, and that performance is stable under label noise and under strongly imbalanced label composition (Figure S7, Table S2). We conducted a one-at-a-time hyperparameter sensitivity analysis over all five loss weights, finding that the total integration score only varies by no more than 0.041 within any sweep across the six scRNA-seq datasets. Thus, the defaults require no per-dataset tuning (Figure S11, Table S5). We explained why cross-platform ST integration is intrinsically hard, showing that as few as 6 of the 2,000 highly variable genes are detected on all six platforms, and quantified where each baseline method fails (Figure 2F, 2G; Figures S20, S21; Tables S6, S7). We further benchmarked FADVI against two spatially-aware methods, GraphST and STAligner (Figures S22; Tables S8). We added a computational efficiency analysis, comparing FADVI with scVI and scANVI (Tables S9). We softened the claim of independent representations, corrected the typos, cited the previously uncited panel, reorganized the supplementary items, and annotated the UMAP figures with quantitative metrics as requested. The FADVI python package was updated to include functionalities for evaluating disentanglement.

## References

1. Luecken, M.D., Gigante, S., Burkhardt, D.B., Cannoodt, R., Strobl, D.C., Markov, N.S., Zappia, L., Palla, G., Lewis, W., Dimitrov, D., et al. (2025). Defining and benchmarking open problems in single-cell analysis. Nat Biotechnol 43, 1035–1040. 10.1038/s41587-025-02694-w.

2. Luecken, M.D., Buttner, M., Chaichoompu, K., Danese, A., Interlandi, M., Mueller, M.F., Strobl, D.C., Zappia, L., Dugas, M., Colome-Tatche, M., and Theis, F.J. (2022). Benchmarking atlas-level data integration in single-cell genomics. Nat Methods 19, 41–50. 10.1038/s41592-021-01336-8.

3. Xu, C., Lopez, R., Mehlman, E., Regier, J., Jordan, M.I., and Yosef, N. (2021). Probabilistic harmonization and annotation of single-cell transcriptomics data with deep generative models. Mol Syst Biol 17, e9620. 10.15252/msb.20209620.

4. Oliveira, M.F., Romero, J.P., Chung, M., Williams, S.R., Gottscho, A.D., Gupta, A., Pilipauskas, S.E., Mohabbat, S., Raman, N., Sukovich, D.J., et al. (2025). High-definition spatial transcriptomic profiling of immune cell populations in colorectal cancer. Nat Genet 57, 1512–1523. 10.1038/s41588-025-02193-3.

5. Stahl, P.L., Salmen, F., Vickovic, S., Lundmark, A., Navarro, J.F., Magnusson, J., Giacomello, S., Asp, M., Westholm, J.O., Huss, M., et al. (2016). Visualization and analysis of gene expression in tissue sections by spatial transcriptomics. Science 353, 78–82. 10.1126/science.aaf2403.

6. He, S., Bhatt, R., Brown, C., Brown, E.A., Buhr, D.L., Chantranuvatana, K., Danaher, P., Dunaway, D., Garrison, R.G., Geiss, G., et al. (2022). High-plex imaging of RNA and proteins at subcellular resolution in fixed tissue by spatial molecular imaging. Nat Biotechnol 40, 1794–1806. 10.1038/s41587-022-01483-z.

7. Janesick, A., Shelansky, R., Gottscho, A.D., Wagner, F., Williams, S.R., Rouault, M., Beliakoff, G., Morrison, C.A., Oliveira, M.F., Sicherman, J.T., et al. (2023). High resolution mapping of the tumor microenvironment using integrated single-cell, spatial and in situ analysis. Nat Commun 14, 8353. 10.1038/s41467-023-43458-x.

8. Liu, R., Qian, K., He, X., and Li, H. (2024). Integration of scRNA-seq data by disentangled representation learning with condition domain adaptation. BMC Bioinformatics 25, 116. 10.1186/s12859-024-05706-9.

9. Piran, Z., Cohen, N., Hoshen, Y., and Nitzan, M. (2024). Disentanglement of single-cell data with biolord. Nat Biotechnol 42, 1678–1683. 10.1038/s41587-023-02079-x.

10. Gao, Y., Dong, K., Shan, C., Li, D., and Liu, Q. (2025). Causal disentanglement for single-cell representations and controllable counterfactual generation. Nat Commun 16, 6775. 10.1038/s41467-025-62008-1.

11. Lopez, R., Regier, J., Cole, M.B., Jordan, M.I., and Yosef, N. (2018). Deep generative modeling for single-cell transcriptomics. Nat Methods 15, 1053–1058. 10.1038/s41592-018-0229-2.

12. Johnson, W.E., Li, C., and Rabinovic, A. (2007). Adjusting batch effects in microarray expression data using empirical Bayes methods. Biostatistics 8, 118–127. 10.1093/biostatistics/kxj037.

13. Korsunsky, I., Millard, N., Fan, J., Slowikowski, K., Zhang, F., Wei, K., Baglaenko, Y., Brenner, M., Loh, P.R., and Raychaudhuri, S. (2019). Fast, sensitive and accurate integration of single-cell data with Harmony. Nat Methods 16, 1289–1296. 10.1038/s41592-019-0619-0.

14. Welch, J.D., Kozareva, V., Ferreira, A., Vanderburg, C., Martin, C., and Macosko, E.Z. (2019). Single-Cell Multi-omic Integration Compares and Contrasts Features of Brain Cell Identity. Cell 177, 1873–1887 e1817. 10.1016/j.cell.2019.05.006.

15. Moinfar, A.A., and Theis, F.J. (2025). Unsupervised Deep Disentangled Representation of Single-Cell Omics. bioRxiv, 2024.2011.2006.622266. 10.1101/2024.11.06.622266.

16. Wilson, P.C., Muto, Y., Wu, H., Karihaloo, A., Waikar, S.S., and Humphreys, B.D. (2022). Multimodal single cell sequencing implicates chromatin accessibility and genetic background in diabetic kidney disease progression. Nat Commun 13, 5253. 10.1038/s41467-022-32972-z.

17. Eraslan, G., Drokhlyansky, E., Anand, S., Fiskin, E., Subramanian, A., Slyper, M., Wang, J., Van Wittenberghe, N., Rouhana, J.M., Waldman, J., et al. (2022). Single-nucleus cross-tissue molecular reference maps toward understanding disease gene function. Science 376, eabl4290. 10.1126/science.abl4290.

18. Steuernagel, L., Lam, B.Y.H., Klemm, P., Dowsett, G.K.C., Bauder, C.A., Tadross, J.A., Hitschfeld, T.S., Del Rio Martin, A., Chen, W., de Solis, A.J., et al. (2022). HypoMap-a unified single-cell gene expression atlas of the murine hypothalamus. Nat Metab 4, 1402–1419. 10.1038/s42255-022-00657-y.

19. Dominguez Conde, C., Xu, C., Jarvis, L.B., Rainbow, D.B., Wells, S.B., Gomes, T., Howlett, S.K., Suchanek, O., Polanski, K., King, H.W., et al. (2022). Cross-tissue immune cell analysis reveals tissue-specific features in humans. Science 376, eabl5197. 10.1126/science.abl5197.

20. Hrovatin, K., Bastidas-Ponce, A., Bakhti, M., Zappia, L., Buttner, M., Salinno, C., Sterr, M., Bottcher, A., Migliorini, A., Lickert, H., and Theis, F.J. (2023). Delineating mouse beta-cell identity during lifetime and in diabetes with a single cell atlas. Nat Metab 5, 1615–1637. 10.1038/s42255-023-00876-x.

21. Tabula Sapiens, C., Jones, R.C., Karkanias, J., Krasnow, M.A., Pisco, A.O., Quake, S.R., Salzman, J., Yosef, N., Bulthaup, B., Brown, P., et al. (2022). The Tabula Sapiens: A multiple-organ, single-cell transcriptomic atlas of humans. Science 376, eabl4896. 10.1126/science.abl4896.

22. Lundberg, S.M., and Lee, S.-I. (2017). A unified approach to interpreting model predictions. Advances in neural information processing systems 30.

23. Ren, P., Zhang, R., Wang, Y., Zhang, P., Luo, C., Wang, S., Li, X., Zhang, Z., Zhao, Y., He, Y., et al. (2025). Systematic benchmarking of high-throughput subcellular spatial transcriptomics platforms across human tumors. Nat Commun 16, 9232. 10.1038/s41467-025-64292-3.

24. Biancalani, T., Scalia, G., Buffoni, L., Avasthi, R., Lu, Z., Sanger, A., Tokcan, N., Vanderburg, C.R., Segerstolpe, A., Zhang, M., et al. (2021). Deep learning and alignment of spatially resolved single-cell transcriptomes with Tangram. Nat Methods 18, 1352–1362. 10.1038/s41592-021-01264-7.

25. Long, Y., Ang, K.S., Li, M., Chong, K.L.K., Sethi, R., Zhong, C., Xu, H., Ong, Z., Sachaphibulkij, K., Chen, A., et al. (2023). Spatially informed clustering, integration, and deconvolution of spatial transcriptomics with GraphST. Nat Commun 14, 1155. 10.1038/s41467-023-36796-3.

26. Zhou, X., Dong, K., and Zhang, S. (2023). Integrating spatial transcriptomics data across different conditions, technologies and developmental stages. Nat Comput Sci 3, 894–906. 10.1038/s43588-023-00528-w.

27. Virshup, I., Bredikhin, D., Heumos, L., Palla, G., Sturm, G., Gayoso, A., Kats, I., Koutrouli, M., Scverse, C., Berger, B., et al. (2023). The scverse project provides a computational ecosystem for single-cell omics data analysis. Nat Biotechnol 41, 604–606. 10.1038/s41587-023-01733-8.

