## Supplementary material for "FADVI: disentangled representation learning for robust integration of single-cell and spatial omics data": Figure S

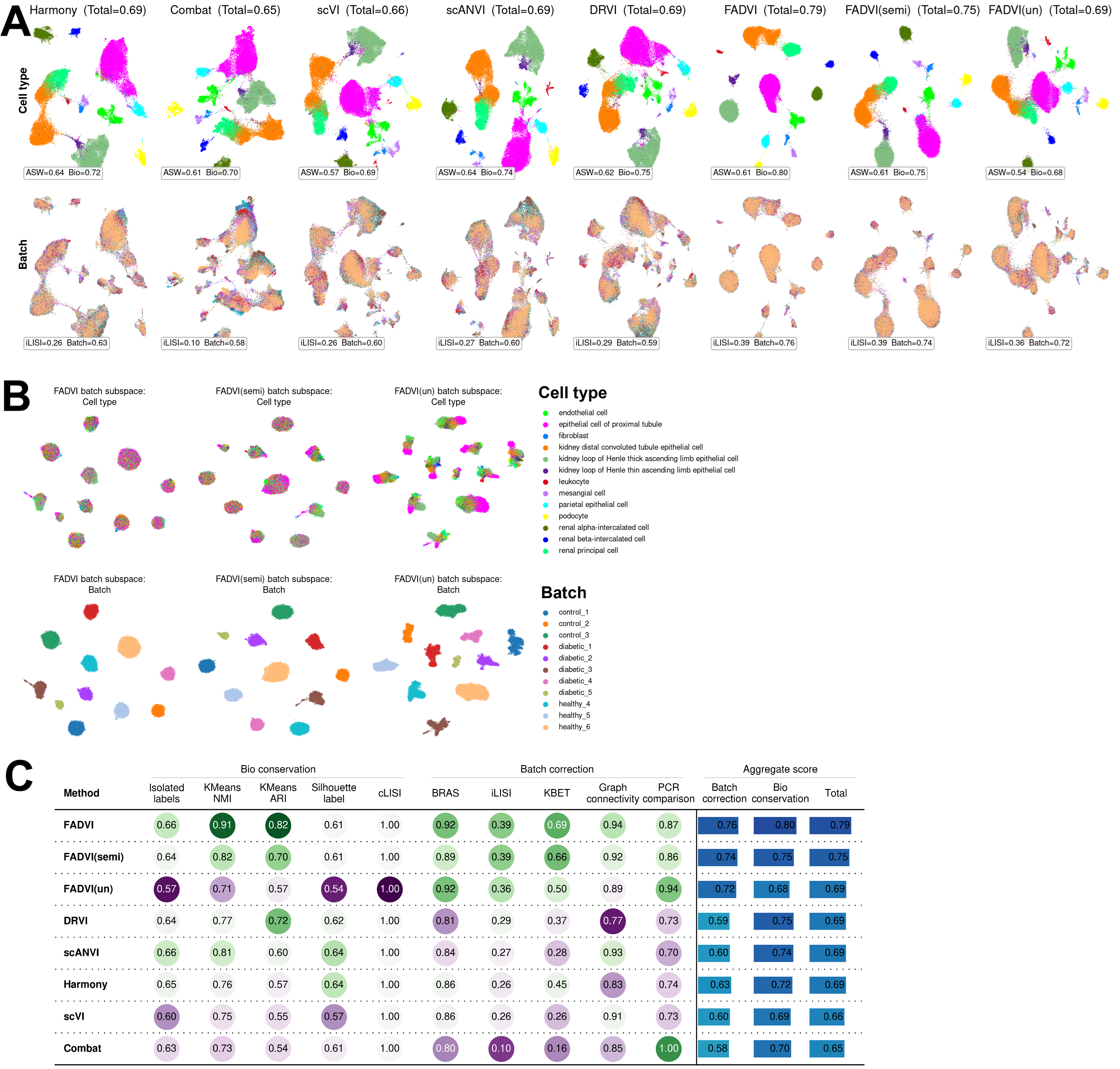

**Figure S1. FADVI demonstrates best data integration in diabetic kidney disease dataset.** (A) UMAP plots of cell type labels and batches using different integration methods. FADVI(semi): semi-supervised training with masked cell type labels in half batches. FADVI(un): unsupervised training without cell type labels. (B) UMAP plots of cell type labels and batches using FADVI representations in batch subspace. (C) Individual and aggregated metrics for evaluating integration methods.

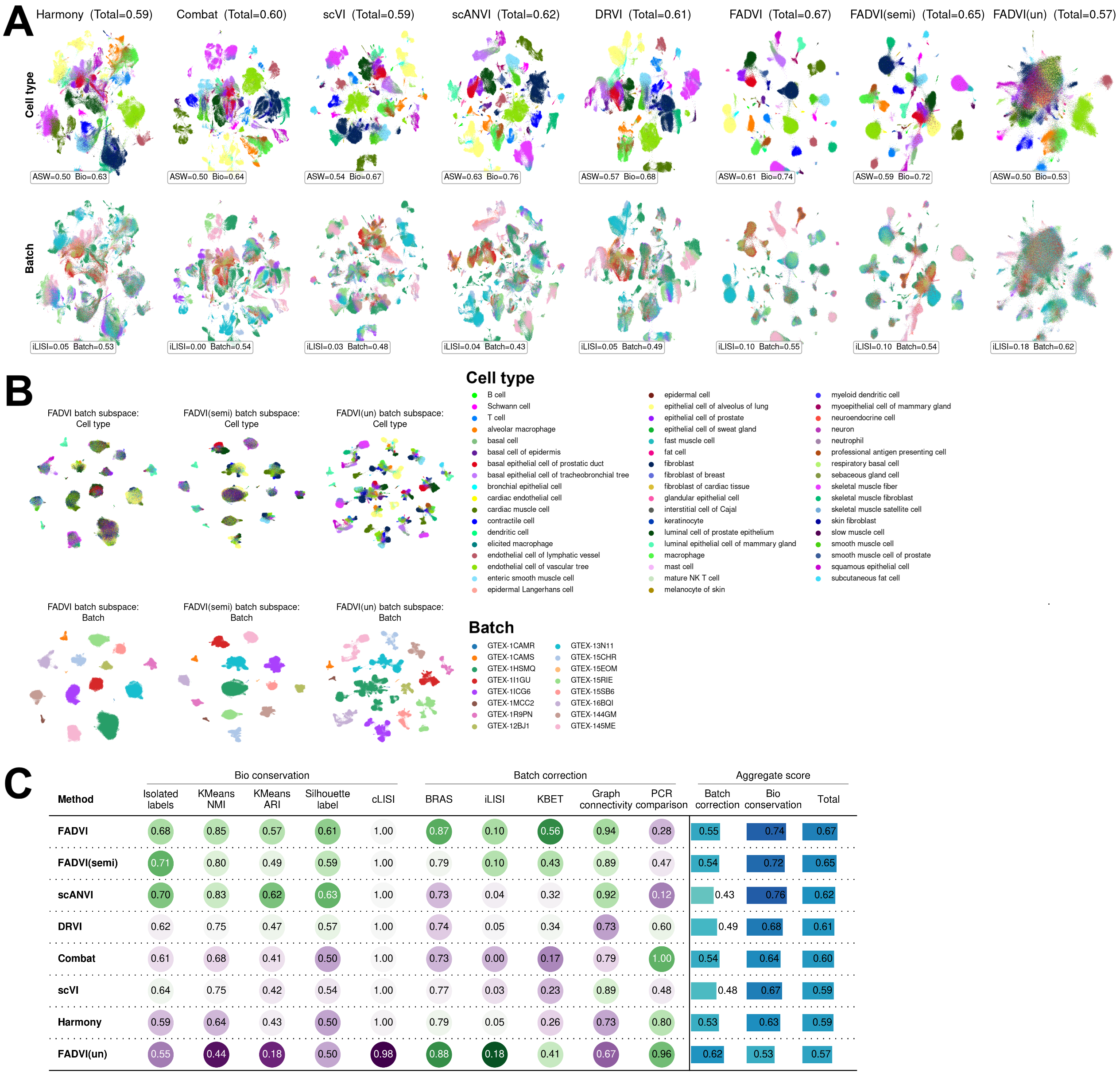

**Figure S2. FADVI demonstrates best data integration in GTEX v9 dataset.** (A) UMAP plots of cell type labels and batches using different integration methods. FADVI(semi): semi-supervised training with masked cell type labels in half batches. FADVI(un): unsupervised training without cell type labels. (B) UMAP plots of cell type labels and batches using FADVI representations in batch subspace. (C) Individual and aggregated metrics for evaluating integration methods.

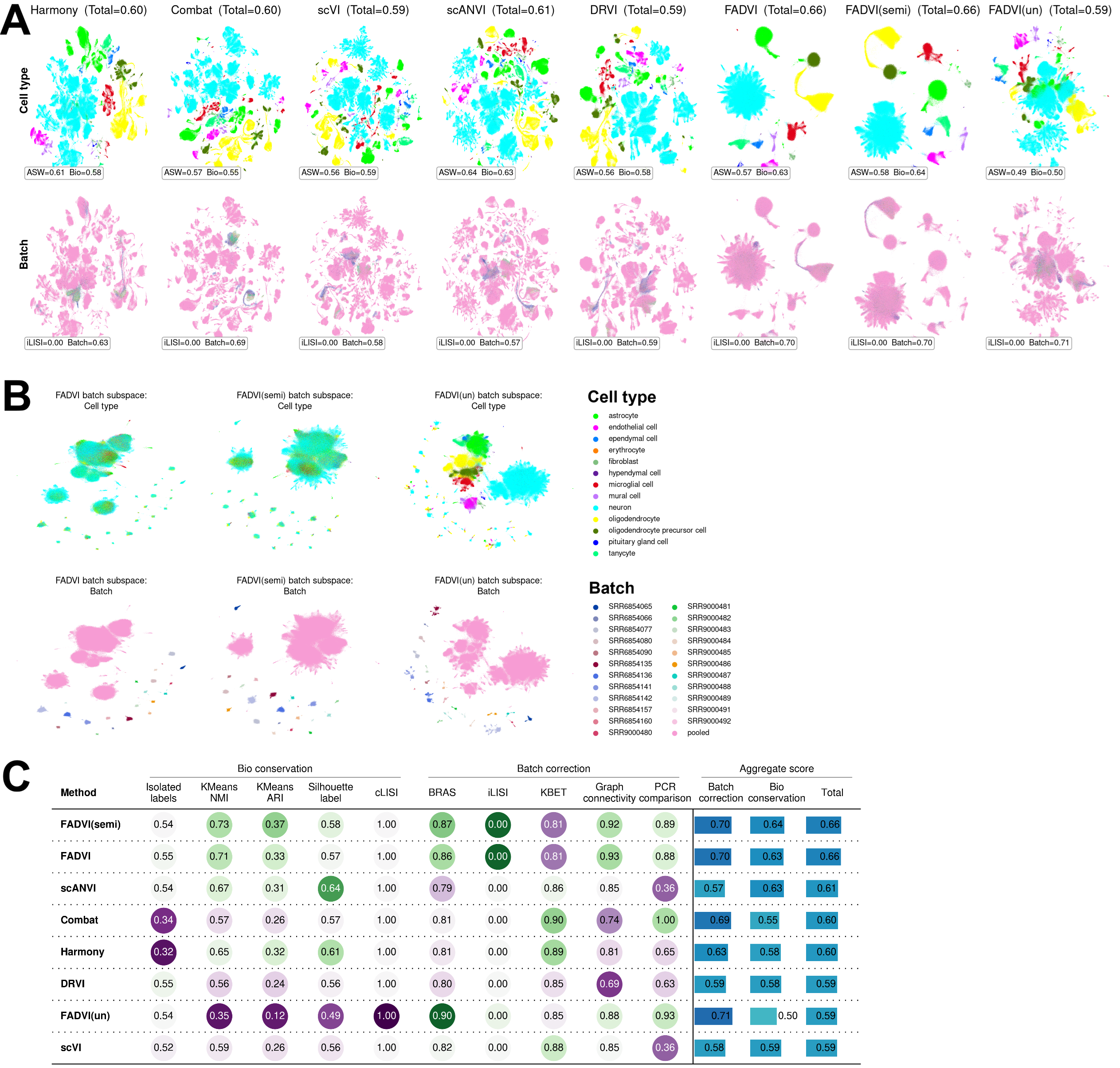

**Figure S3. FADVI demonstrates best data integration in HypoMap dataset.** (A) UMAP plots of cell type labels and batches using different integration methods. FADVI(semi): semi-supervised training with masked cell type labels in half batches. FADVI(un): unsupervised training without cell type labels. (B) UMAP plots of cell type labels and batches using FADVI representations in batch subspace. (C) Individual and aggregated metrics for evaluating integration methods.

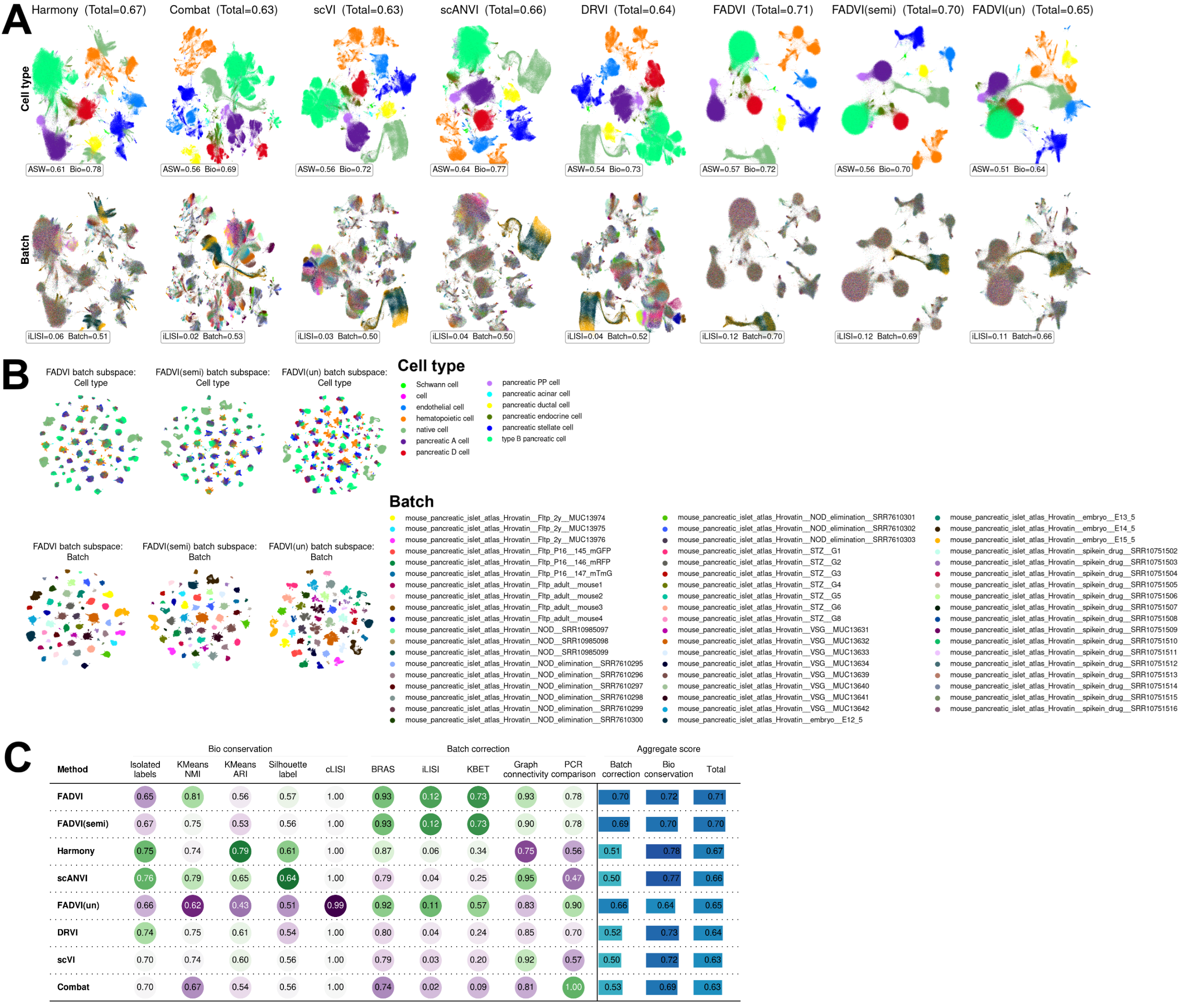

**Figure S4. FADVI demonstrates best data integration in Mouse Pancreatic Islet dataset.** (A) UMAP plots of cell type labels and batches using different integration methods. FADVI(semi): semi-supervised training with masked cell type labels in half batches. FADVI(un): unsupervised training without cell type labels. (B) UMAP plots of cell type labels and batches using FADVI representations in batch subspace. (C) Individual and aggregated metrics for evaluating integration methods.

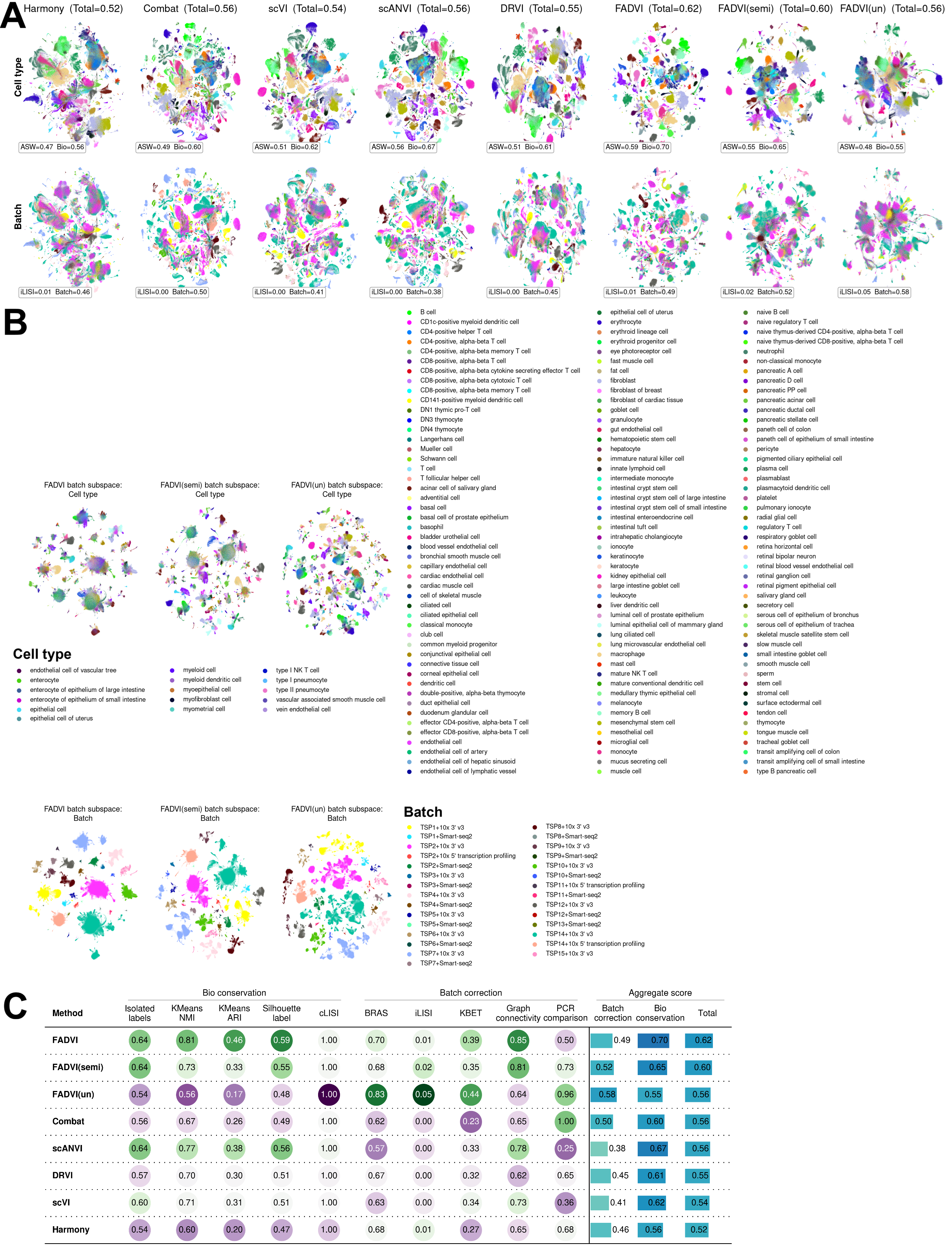

**Figure S5. FADVI demonstrates best data integration in Tabula Sapiens.** (A) UMAP plots of cell type labels and batches using different integration methods. FADVI(semi): semi-supervised training with masked cell type labels in half batches. FADVI(un): unsupervised training without cell type labels. (B) UMAP plots of cell type labels and batches using FADVI representations in batch subspace. (C) Individual and aggregated metrics for evaluating integration methods.

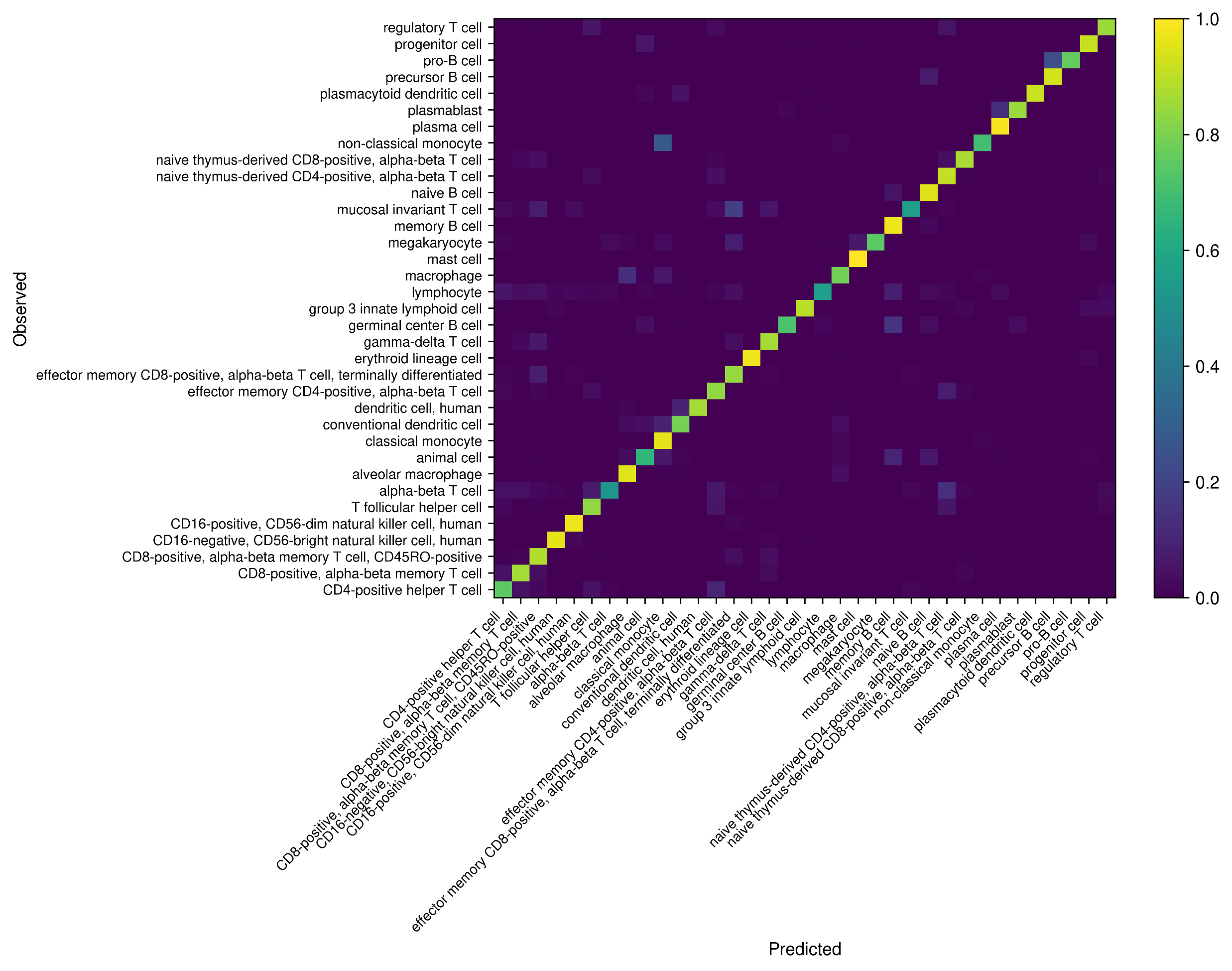

**Figure S6. Semi-supervised FADVI enables accurate label projection in Immune Cell Atlas.** The heatmap shows the ratio of predicted cell type labels for unlabeled cells. FADVI makes accurate prediction for most cells, and the wrong predictions are mainly similar immune cell subtypes such as different T cell subtypes.

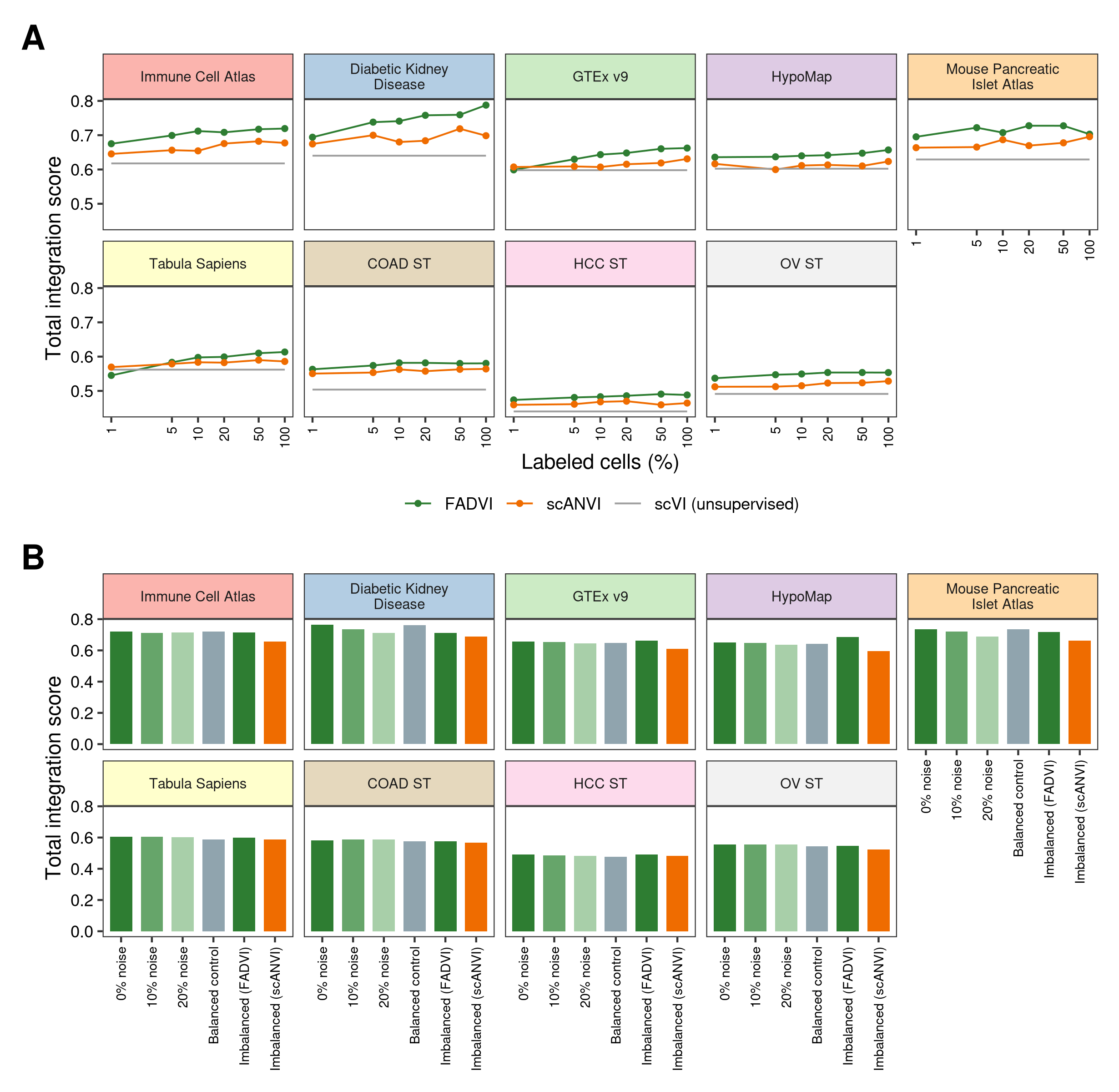

**Figure S7. FADVI maintains integration performance with sparse, noisy, or imbalanced cell type labels.** (A) Total integration score with different percentage of labeled cells (1-100%, stratified within each batch) for FADVI and scANVI across nine datasets. scVI receives no labels and is shown as a constant reference. FADVI retains 88-99% of its fully-supervised performance with only 1% of cells labeled, and exceeds scANVI at nearly every labeled fraction. (B) Total integration score under label corruption and label imbalance. Label noise: the stated fraction of retained labels was randomly permuted. Imbalanced: abundant cell types were labeled at 1% while the remaining types were labeled at 50%, so that labeled and unlabeled subsets differ markedly in composition; the balanced control uses the same total number of labels drawn proportionally. Performance is stable across all conditions.

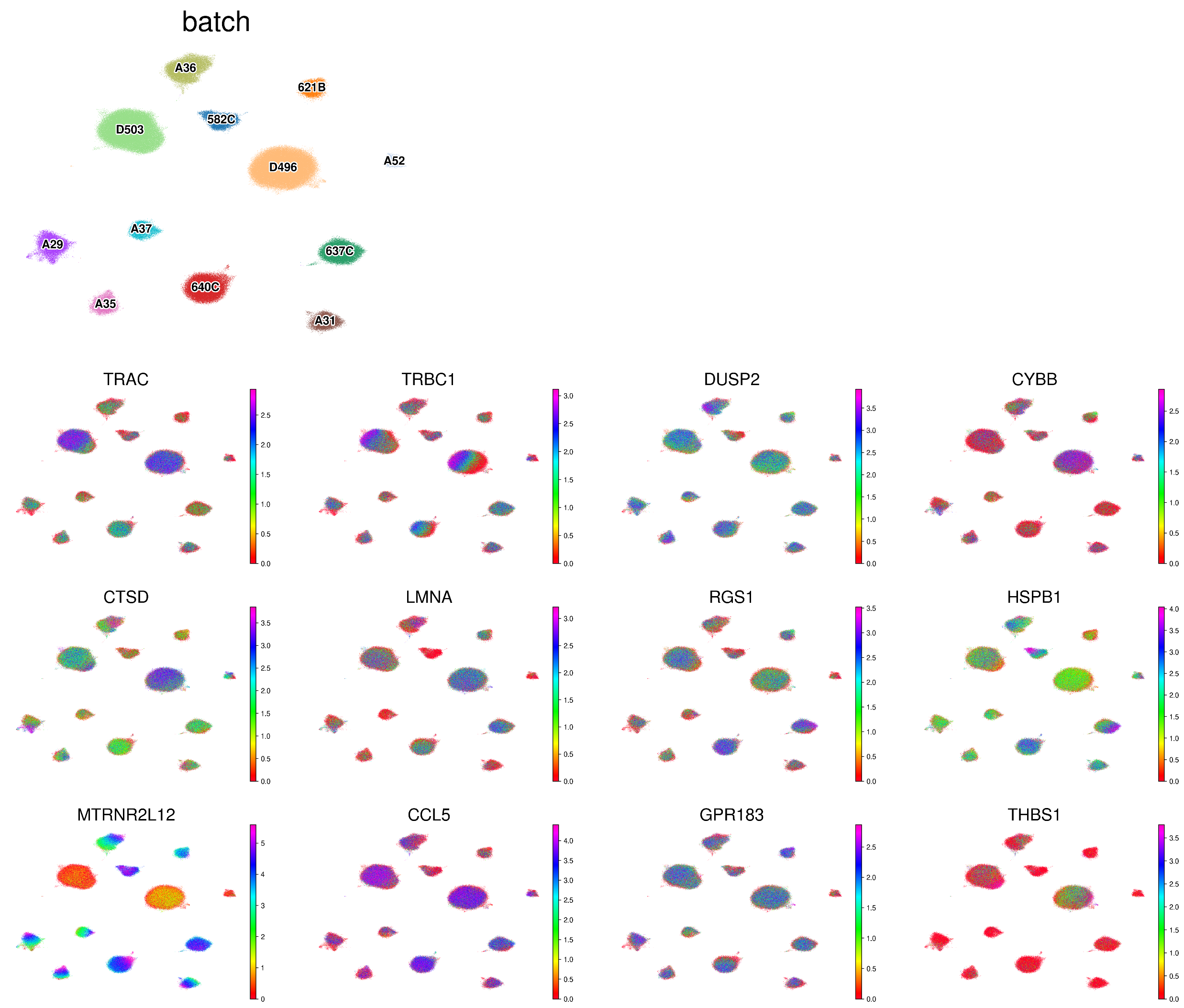

**Figure S8. FADVI identifies representative genes associated with batch effect in Immune Cell Atlas.** UMAP plots of all cells in Immune Cell Atlas show the expression variation across batches in FADVI batch subspace. Genes with high attribution to batches in Figure 1H were selected.

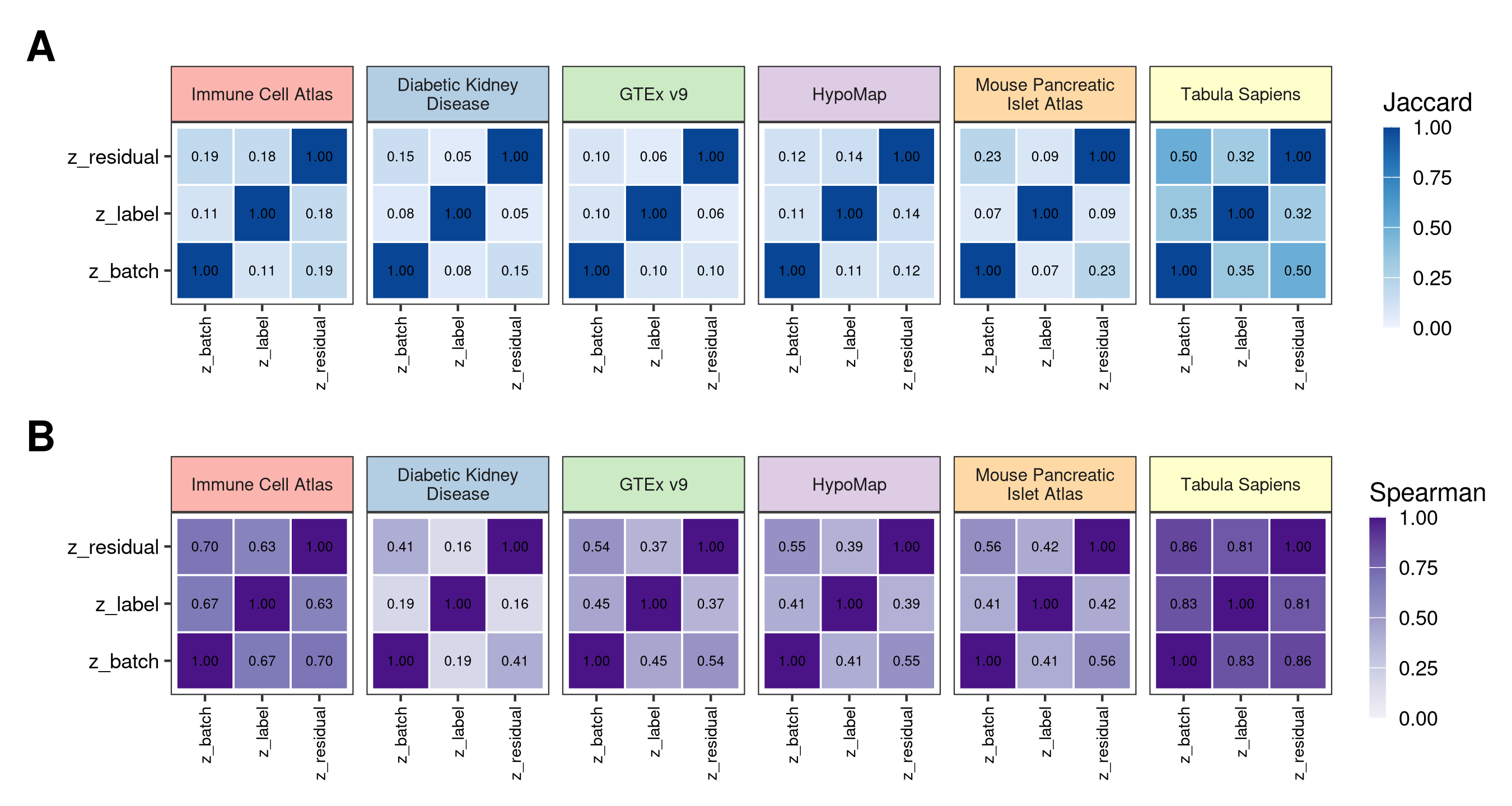

**Figure S9. Feature attribution confirms that the three latent subspaces are driven by distinct genes.** (A) Pairwise Jaccard overlap between the sets of top-attributed genes for the batch (z_batch), label (z_label) and residual (z_residual) subspaces across six scRNA-seq datasets. Low off-diagonal values indicate that each subspace is driven by a largely distinct set of genes. (B) Spearman rank correlation of per-gene attributions between subspaces. Attributions were computed with integrated gradients applied to the L2 norm of each subspace, which allows attribution to the unsupervised residual subspace, and were residualized against expression rank before comparison because raw attribution magnitude correlates strongly with expression level.

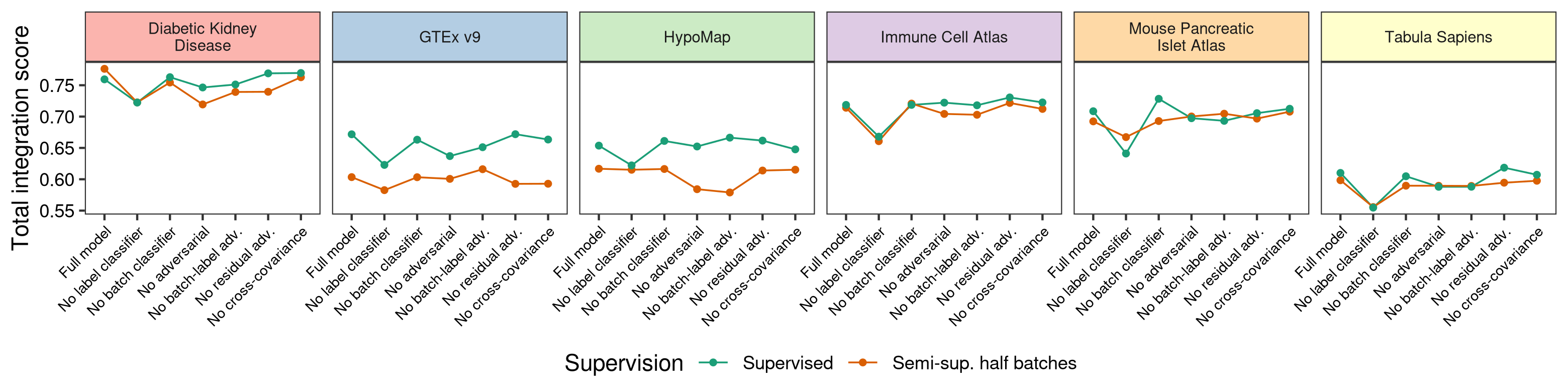

**Figure S10. Label classification loss contributes most to FADVI integration performance.** Total integration score across ablation settings in six scRNA-seq datasets under supervised training and semi-supervised training with masked cell type labels in half of the batches. Each ablation removes one loss term, equivalent to setting its weight to zero; see Figure S11 for the full sweep of loss weights. Removing the label classifier causes the largest decline, while removal of the residual adversarial terms has minimal effect.

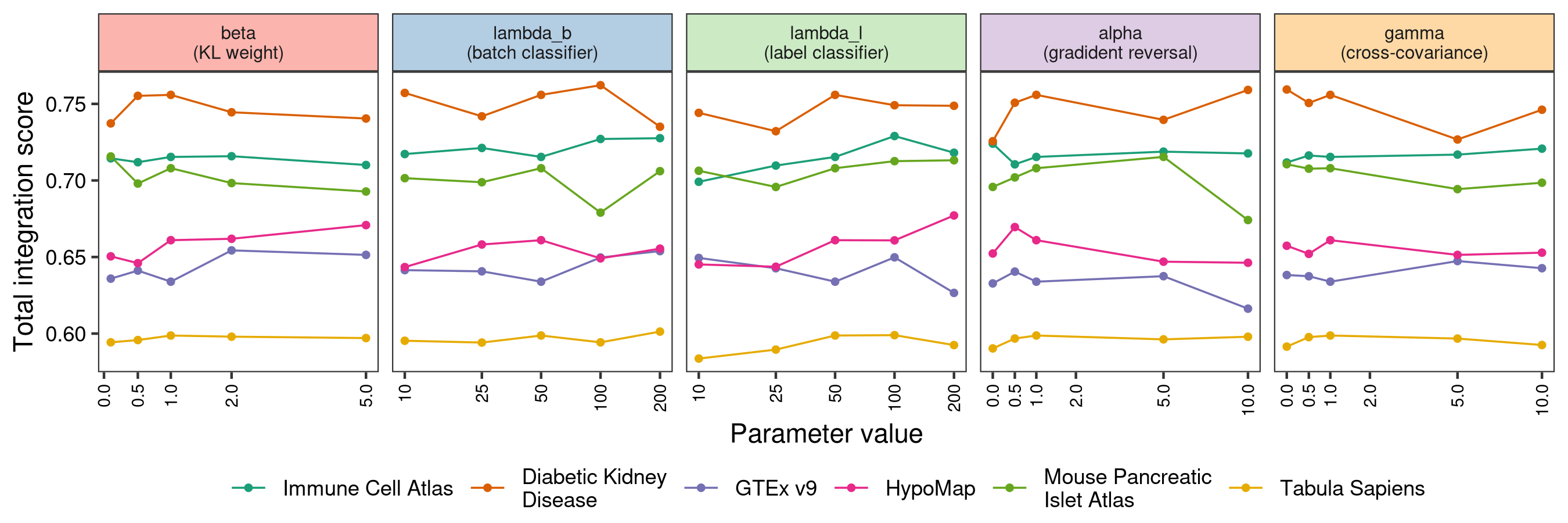

**Figure S11. FADVI integration performance is robust to loss weight hyperparameters.** Total integration score as each loss weight is varied one at a time from the default configuration, with all other weights held at their default values, in six scRNA-seq datasets. All runs are under semi-supervised training with masked cell type labels in half of the batches. beta weights the KL regularization strength, lambda_b and lambda_l weight the batch and label classification losses, alpha weights the gradient reversal strength of the adversarial heads, and gamma weights the cross-covariance penalty. The x-axis uses a pseudo-logarithmic scale so that alpha = 0 and gamma = 0, which switch the adversarial and decorrelation terms off entirely, can be shown. The total score varies by at most 0.041 within any single sweep, indicating that the default settings are near-optimal and that per-dataset tuning is not required.

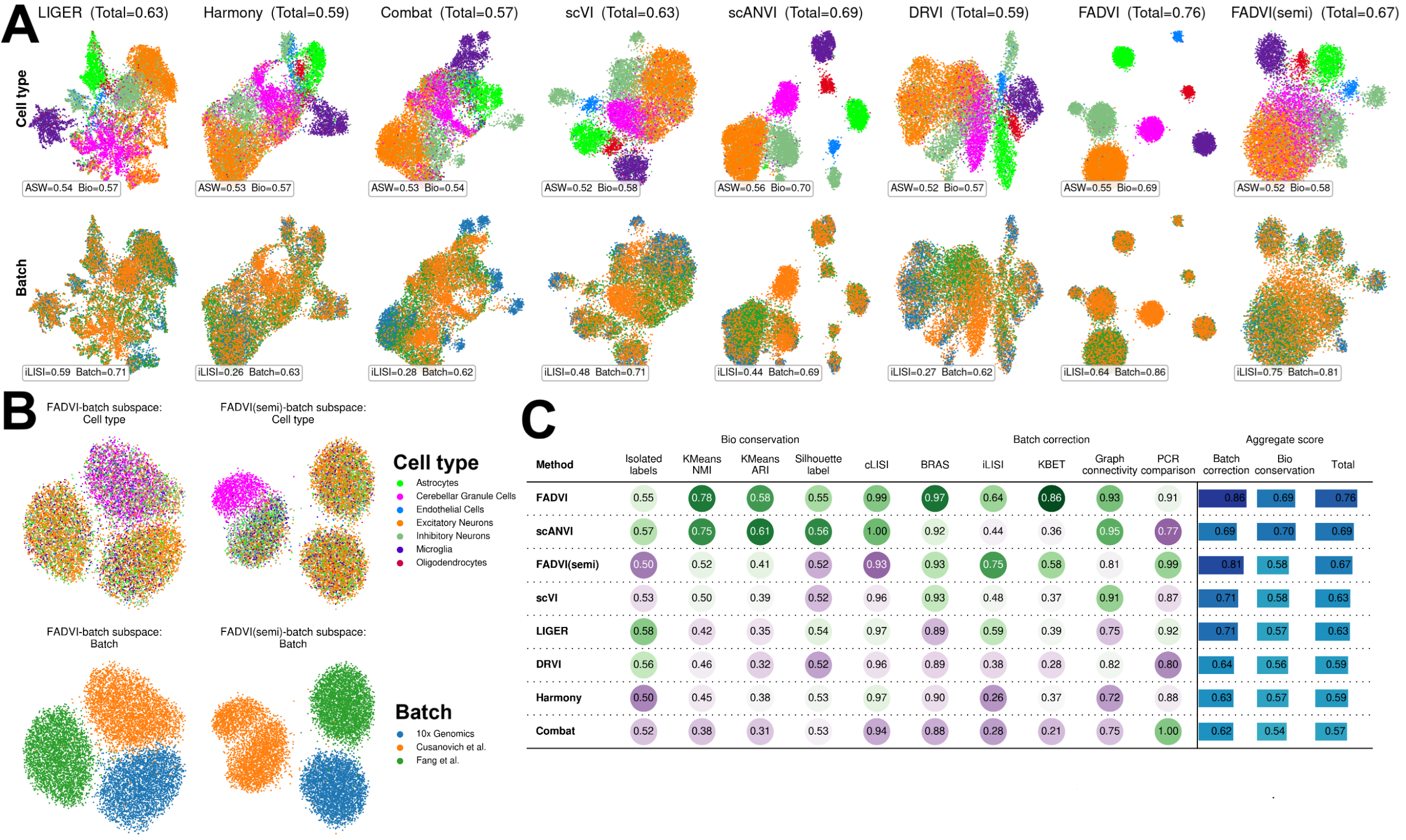

**Figure S12. FADVI demonstrates best data integration in small scATAC-seq dataset with gene activity features.** (A) UMAP plots of cell type labels and batches using different integration methods. FADVI(semi): semi-supervised training with masked cell type labels in two smaller batches. (B) UMAP plots of cell type labels and batches using FADVI representations in batch subspace. (C) Individual and aggregated metrics for evaluating integration methods.

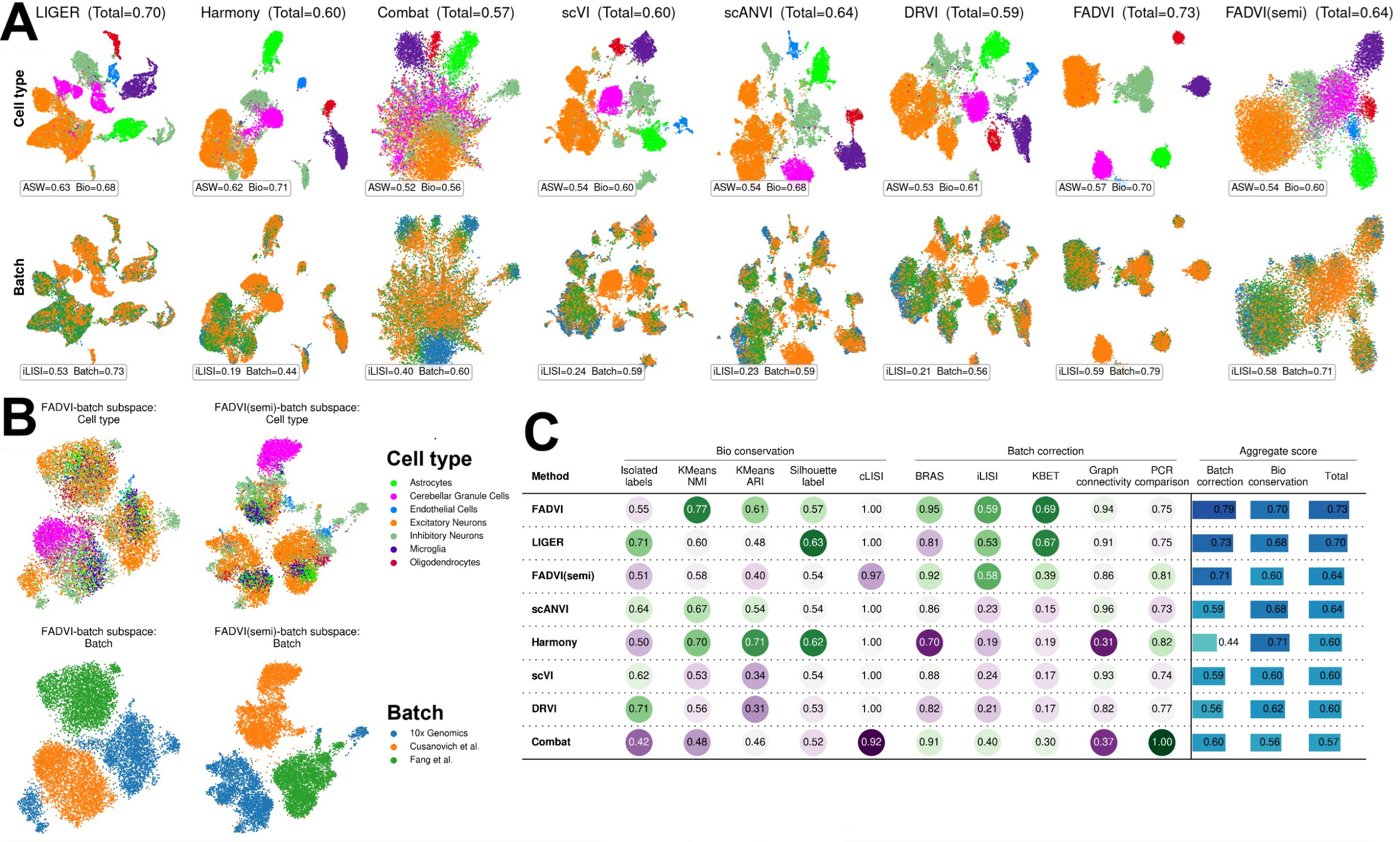

**Figure S13. FADVI demonstrates best data integration in small scATAC-seq dataset with peak features.** (A) UMAP plots of cell type labels and batches using different integration methods. FADVI(semi): semi-supervised training with masked cell type labels in two smaller batches. (B) UMAP plots of cell type labels and batches using FADVI representations in batch subspace. (C) Individual and aggregated metrics for evaluating integration methods.

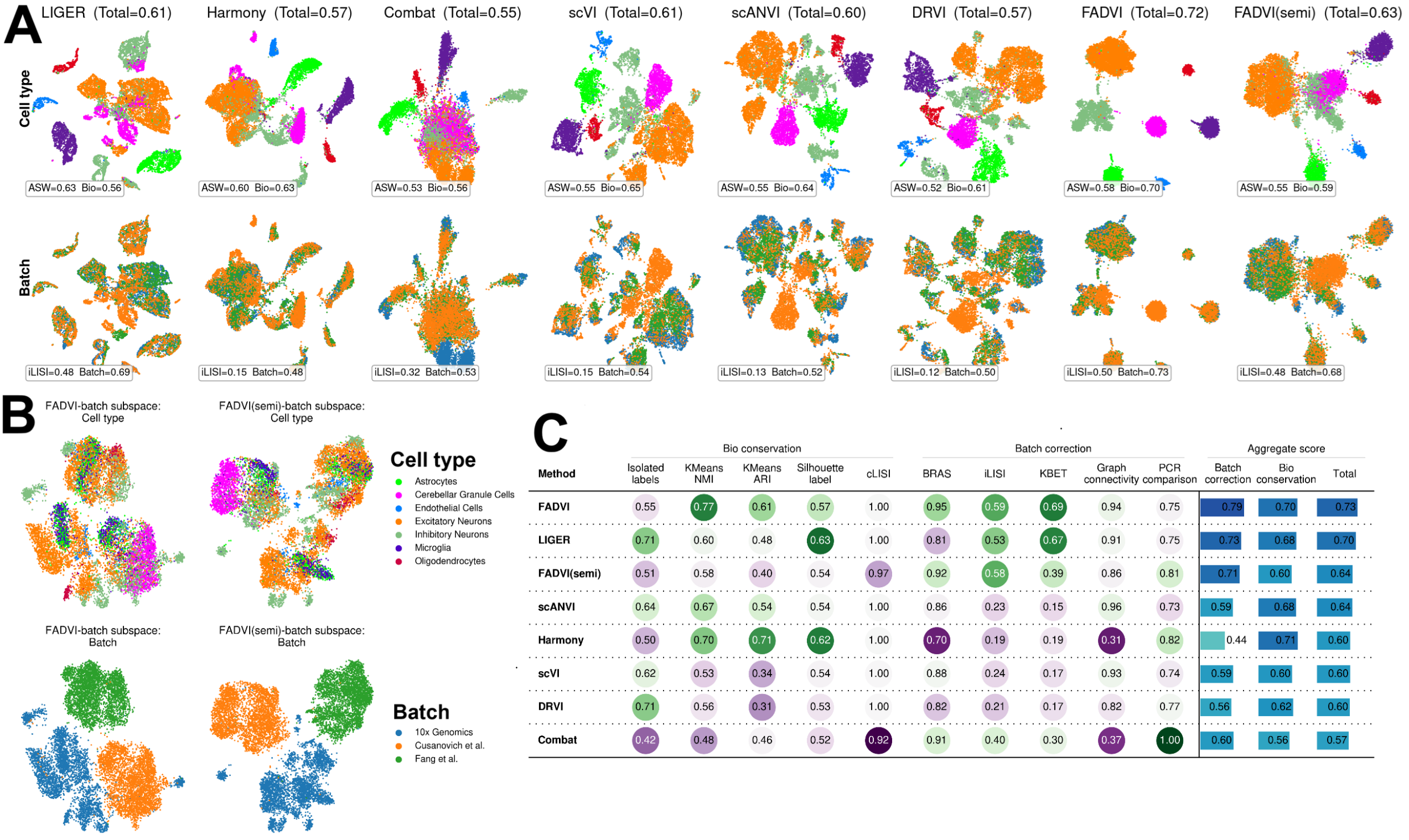

**Figure S14. FADVI demonstrates best data integration in small scATAC-seq dataset with window features.** (A) UMAP plots of cell type labels and batches using different integration methods. FADVI(semi): semi-supervised training with masked cell type labels in two smaller batches. (B) UMAP plots of cell type labels and batches using FADVI representations in batch subspace. (C) Individual and aggregated metrics for evaluating integration methods.

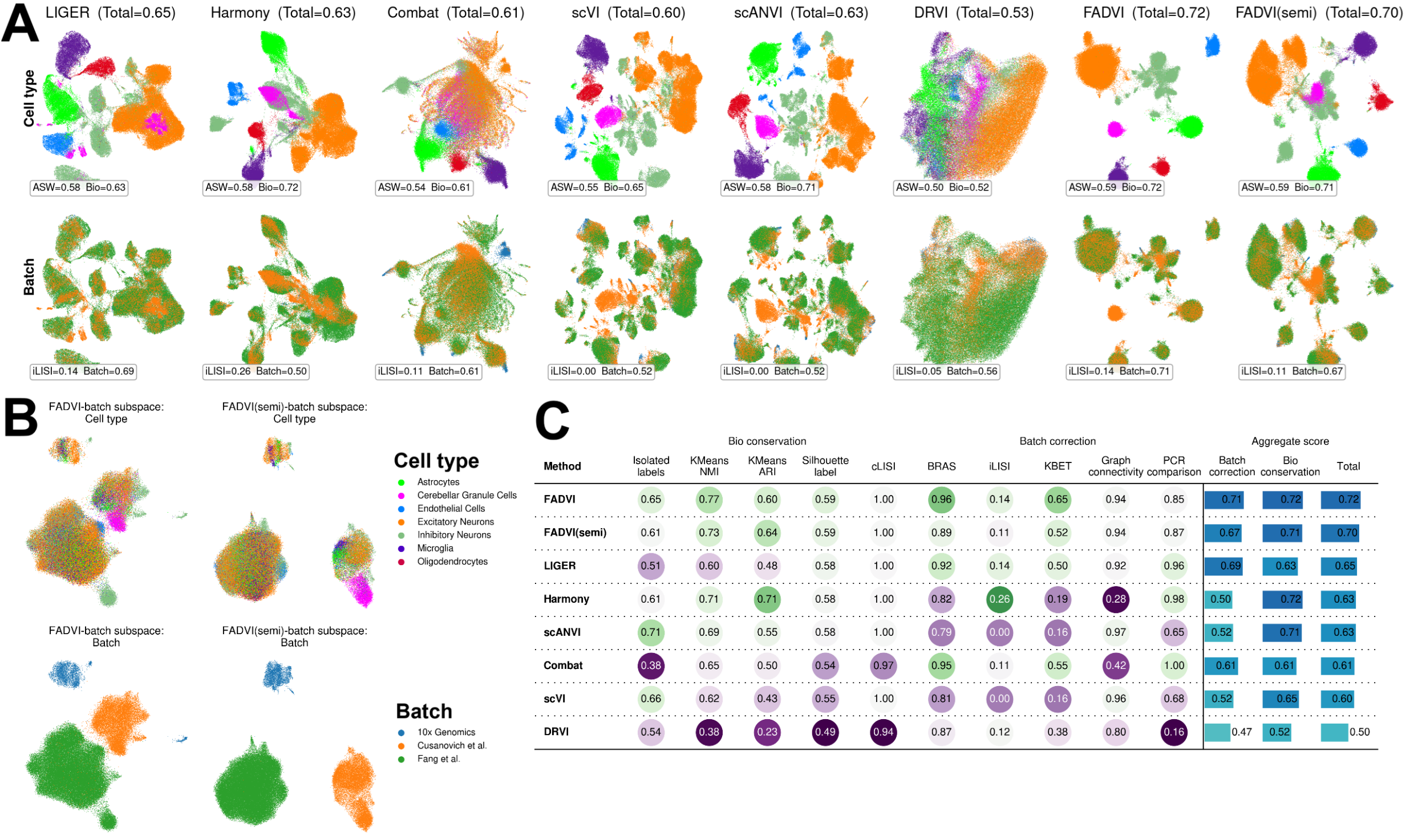

**Figure S15. FADVI demonstrates best data integration in large scATAC-seq dataset with gene activity features.** (A) UMAP plots of cell type labels and batches using different integration methods. FADVI(semi): semi-supervised training with masked cell type labels in two smaller batches. (B) UMAP plots of cell type labels and batches using FADVI representations in batch subspace. (C) Individual and aggregated metrics for evaluating integration methods.

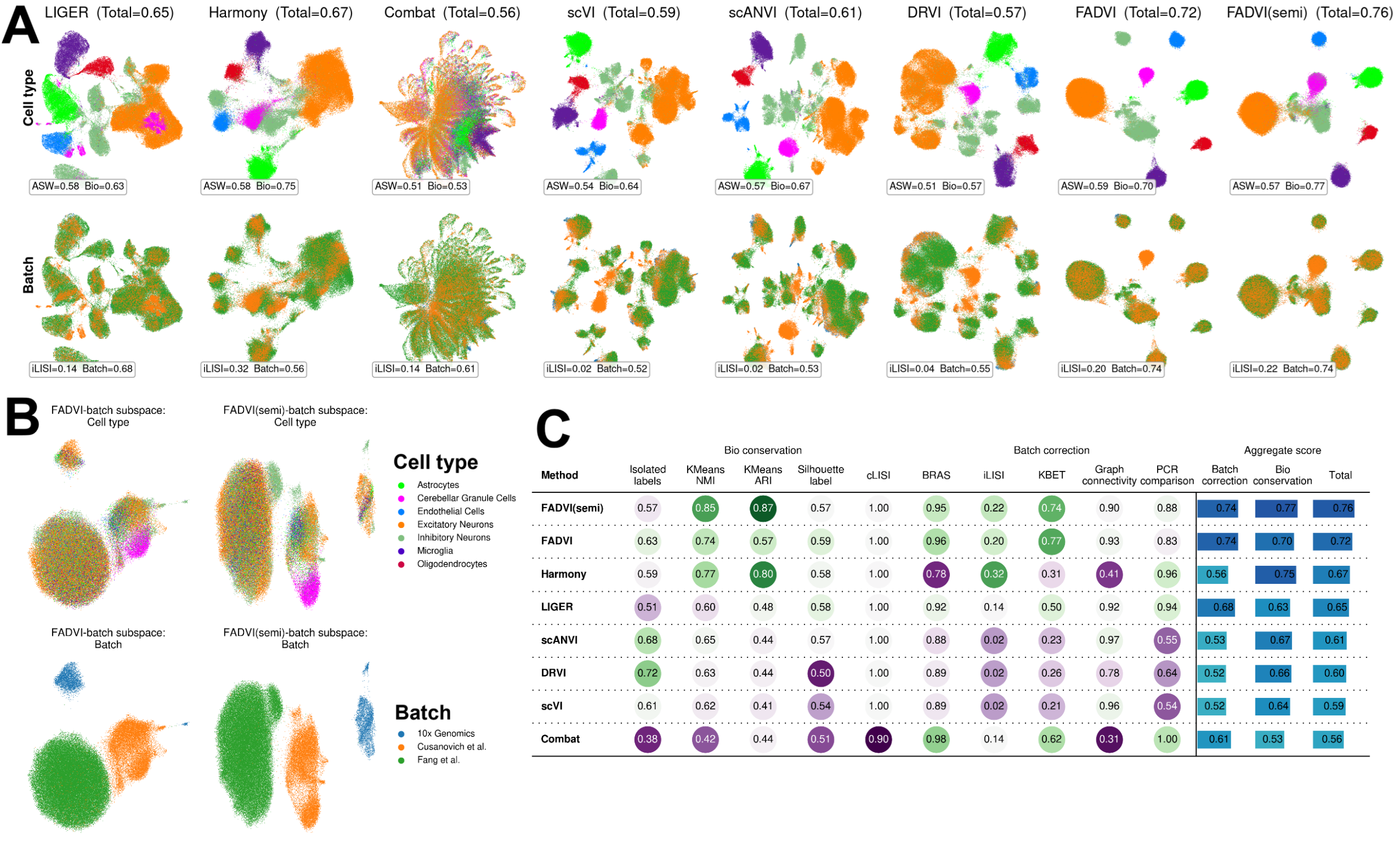

**Figure S16. FADVI demonstrates best data integration in large scATAC-seq dataset with peak features.** (A) UMAP plots of cell type labels and batches using different integration methods. FADVI(semi): semi-supervised training with masked cell type labels in two smaller batches. (B) UMAP plots of cell type labels and batches using FADVI representations in batch subspace. (C) Individual and aggregated metrics for evaluating integration methods.

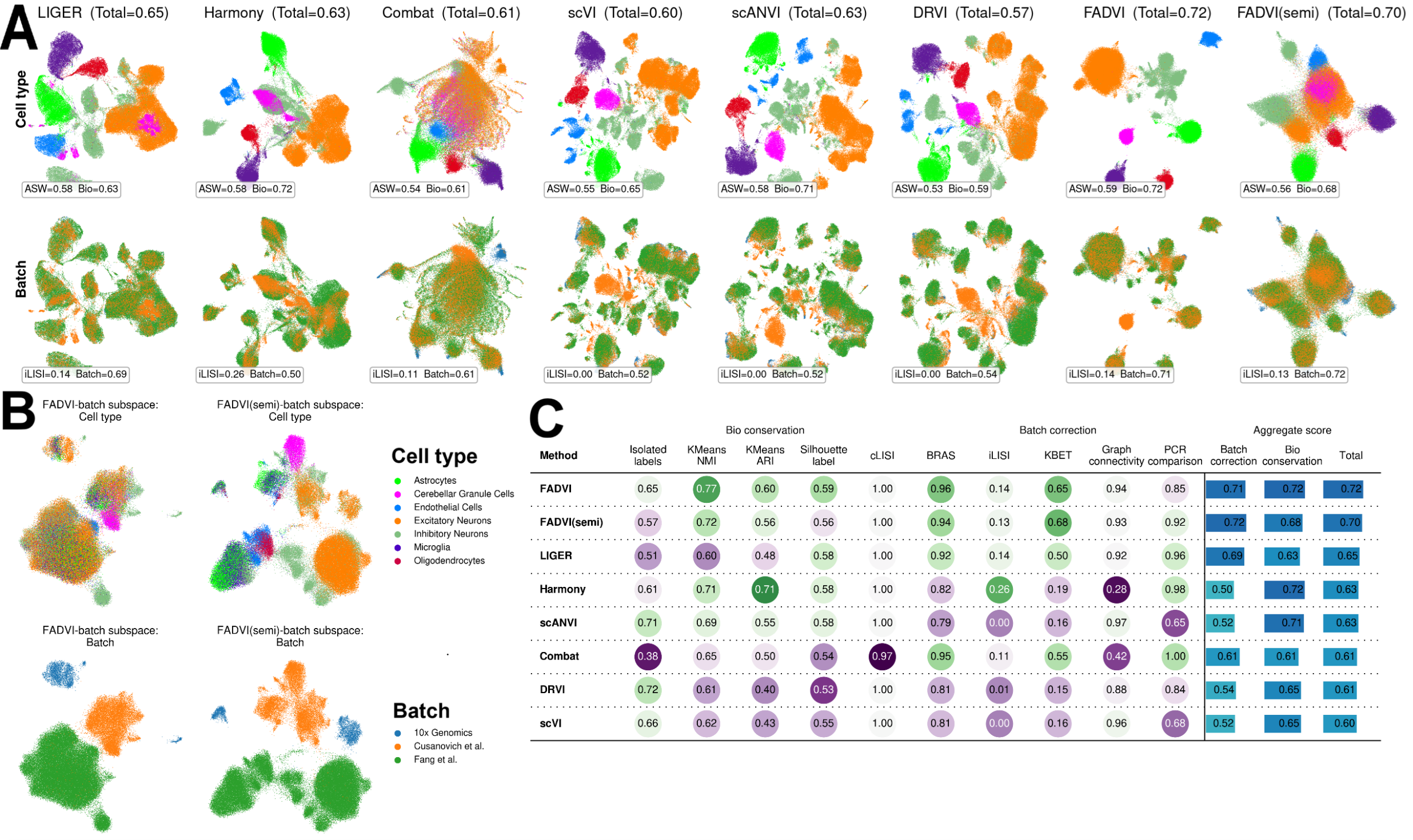

**Figure S17. FADVI demonstrates best data integration in large scATAC-seq dataset with window features.** (A) UMAP plots of cell type labels and batches using different integration methods. FADVI(semi): semi-supervised training with masked cell type labels in two smaller batches. (B) UMAP plots of cell type labels and batches using FADVI representations in batch subspace. (C) Individual and aggregated metrics for evaluating integration methods.

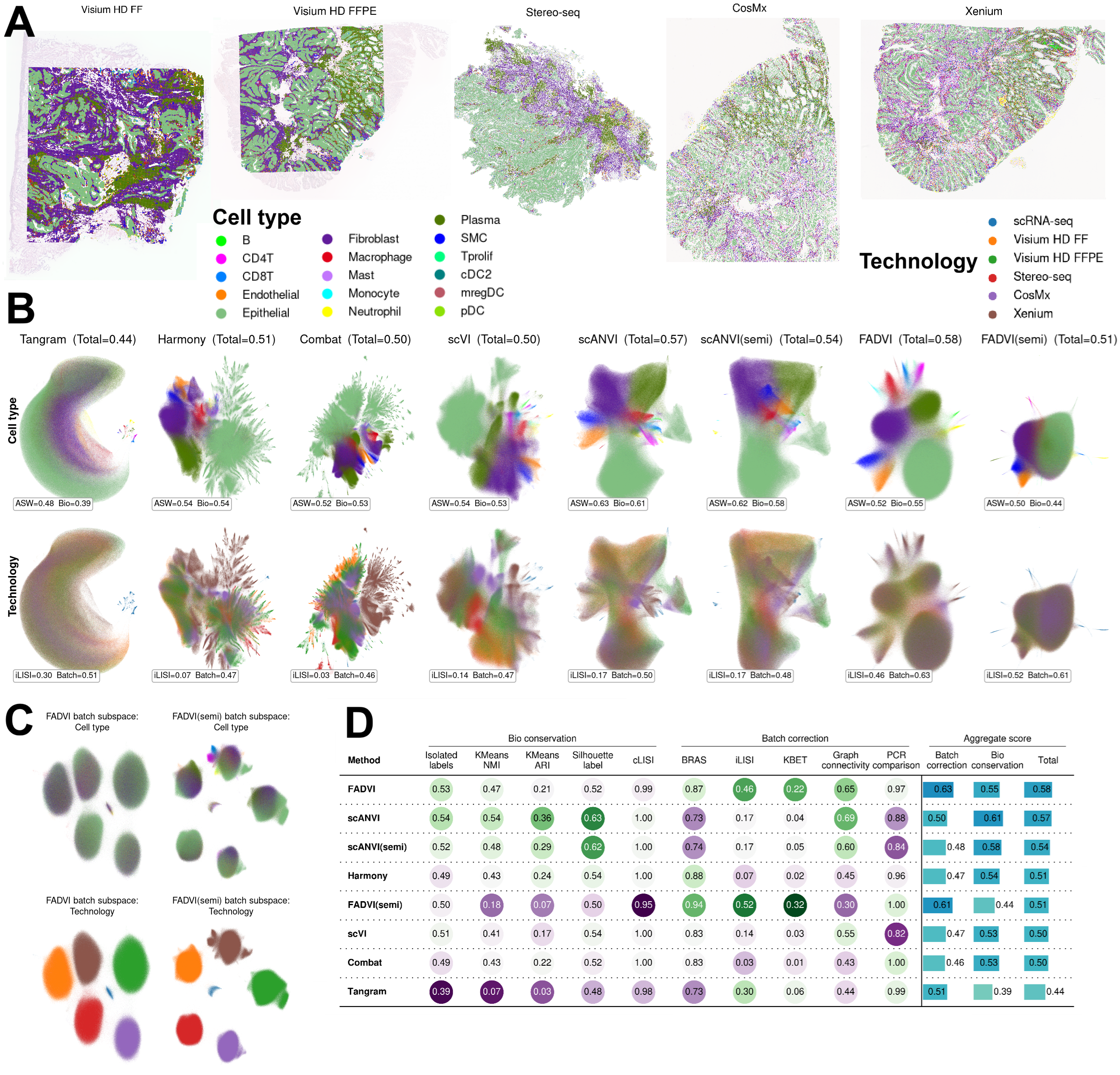

**Figure S18. FADVI demonstrates best data integration in COAD ST with scRNA-seq dataset.** (A) Cell types labels of COAD ST data overlaid on hematoxylin and eosin staining image. (B) UMAP plots of cell type labels and batches using different integration methods. FADVI(semi): semi-supervised training with masked cell type labels in all ST data. (C) UMAP plots of cell type labels and batches using FADVI representations in batch subspace. (D) Individual and aggregated metrics for evaluating integration methods.

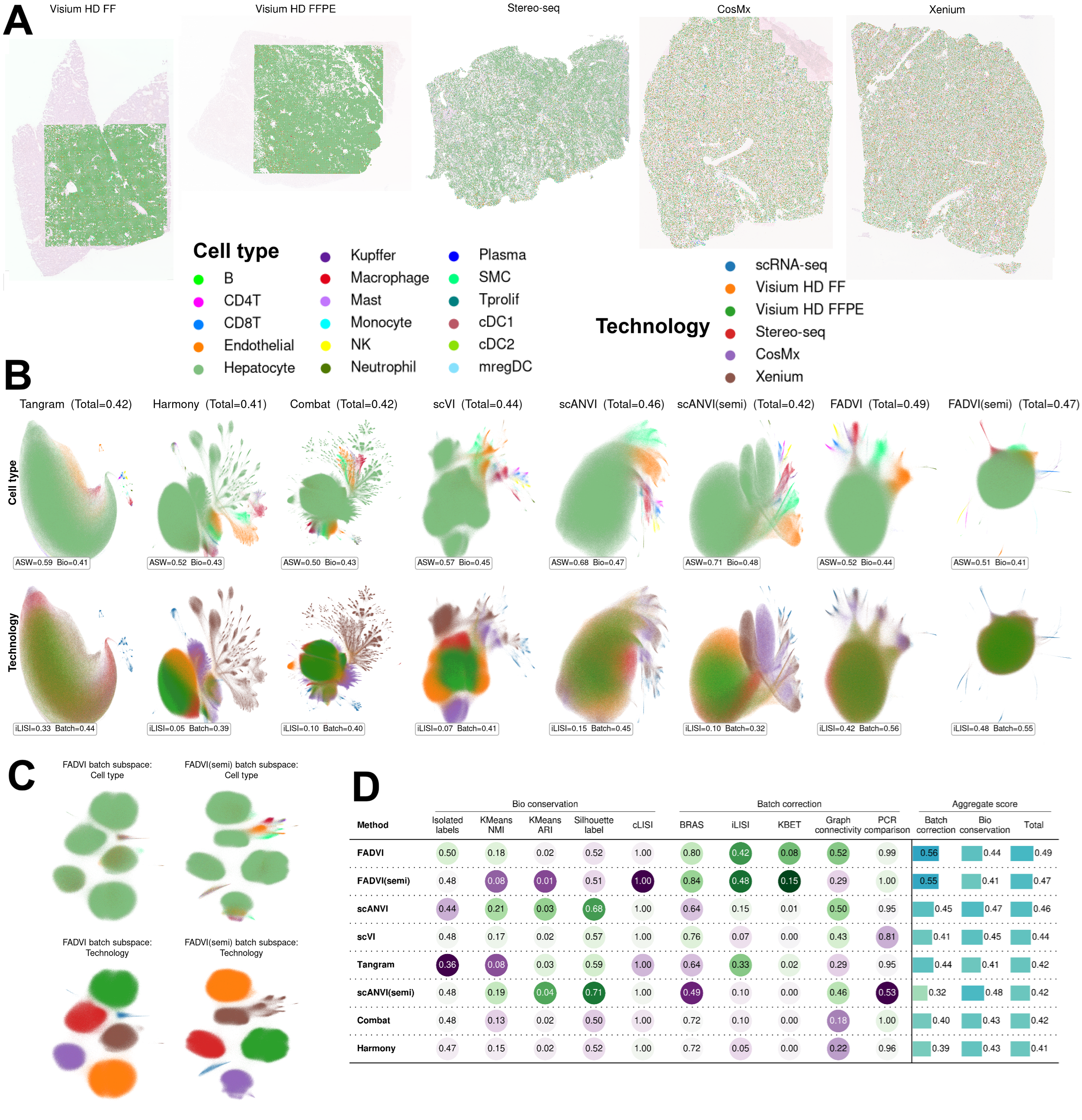

**Figure S19. FADVI demonstrates best data integration HCC ST with scRNA-seq dataset.** (A) Cell types labels of HCC ST data overlaid on hematoxylin and eosin staining image. (B) UMAP plots of cell type labels and batches using different integration methods. FADVI(semi): semi-supervised training with masked cell type labels in all ST data. (C) UMAP plots of cell type labels and batches using FADVI representations in batch subspace. (D) Individual and aggregated metrics for evaluating integration methods.

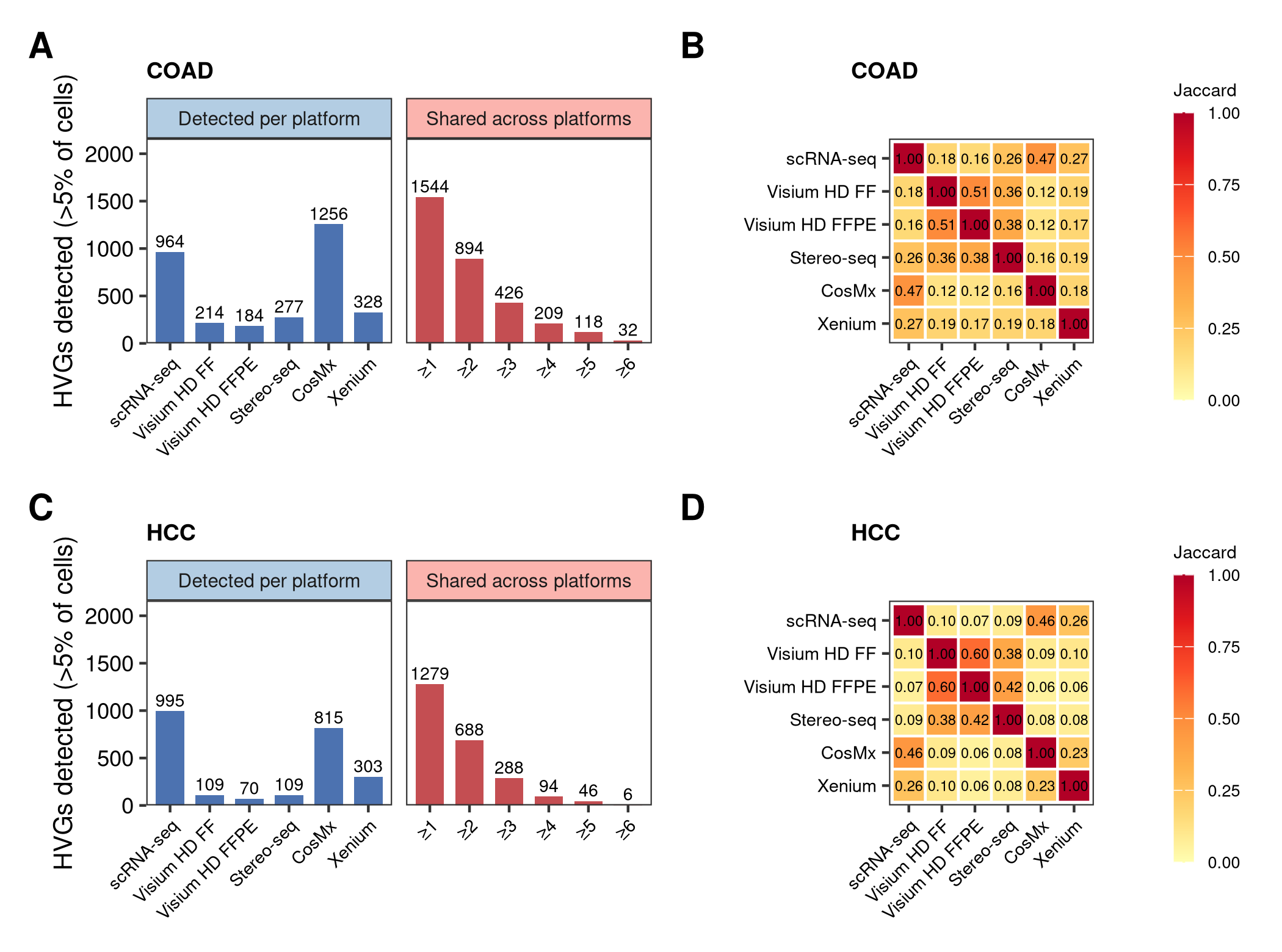

**Figure S20. Highly variable genes are largely non-overlapping across spatial platforms in COAD and HCC ST datasets.** (A, C) Number of the 2,000 highly variable genes detected in more than 5% of cells on each platform, and the number detected on at least k platforms (k = 1-6), for COAD (A) and HCC (C). Only 32 genes (1.6%) in COAD and 6 genes (0.3%) in HCC are detected on all six platforms. (B, D) Pairwise Jaccard overlap between the detected-gene sets of each platform pair for COAD (B) and HCC (D). Overlap is lowest between imaging-based and sequencing-based platforms. Because the shared feature space is nearly empty, methods that assume a common feature distribution have little to align on. Corresponding results for the OV dataset are shown in Figure 2F and 2G.

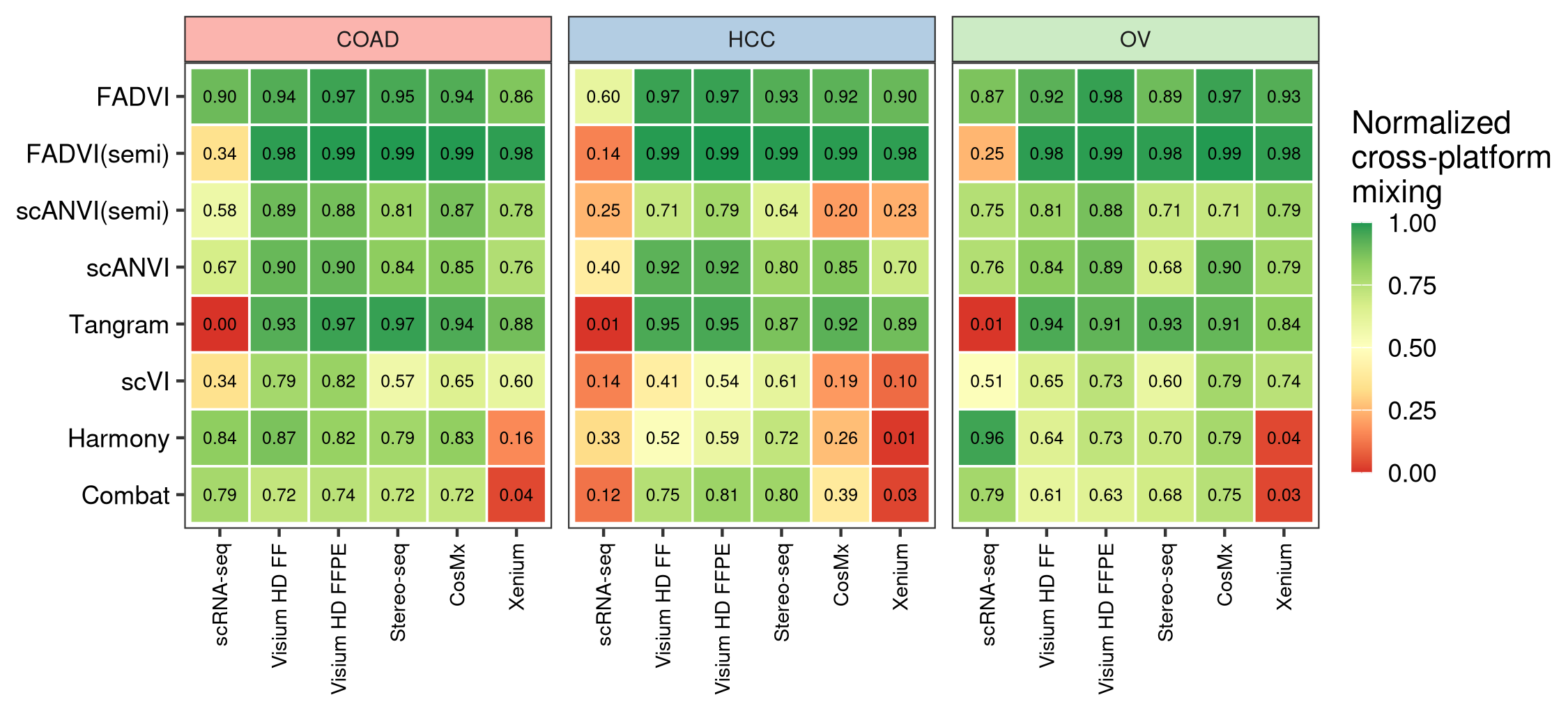

**Figure S21. Baseline integration methods fail on specific spatial platforms while FADVI does not.** Normalized cross-platform mixing for each integration method and platform in the three ST datasets, defined as the fraction of a cell’s k=30 nearest neighbors drawn from other platforms, normalized by the size-based null expectation (1 = fully integrated, 0 = an isolated island of cells). Values below 0.5 indicate failure to integrate that platform. Xenium is the most common point of failure, and FADVI is the only method that never fails on any platform-dataset combination. The low scRNA-seq value for Tangram is expected by design, as Tangram maps ST data onto the scRNA-seq reference rather than co-embedding both.

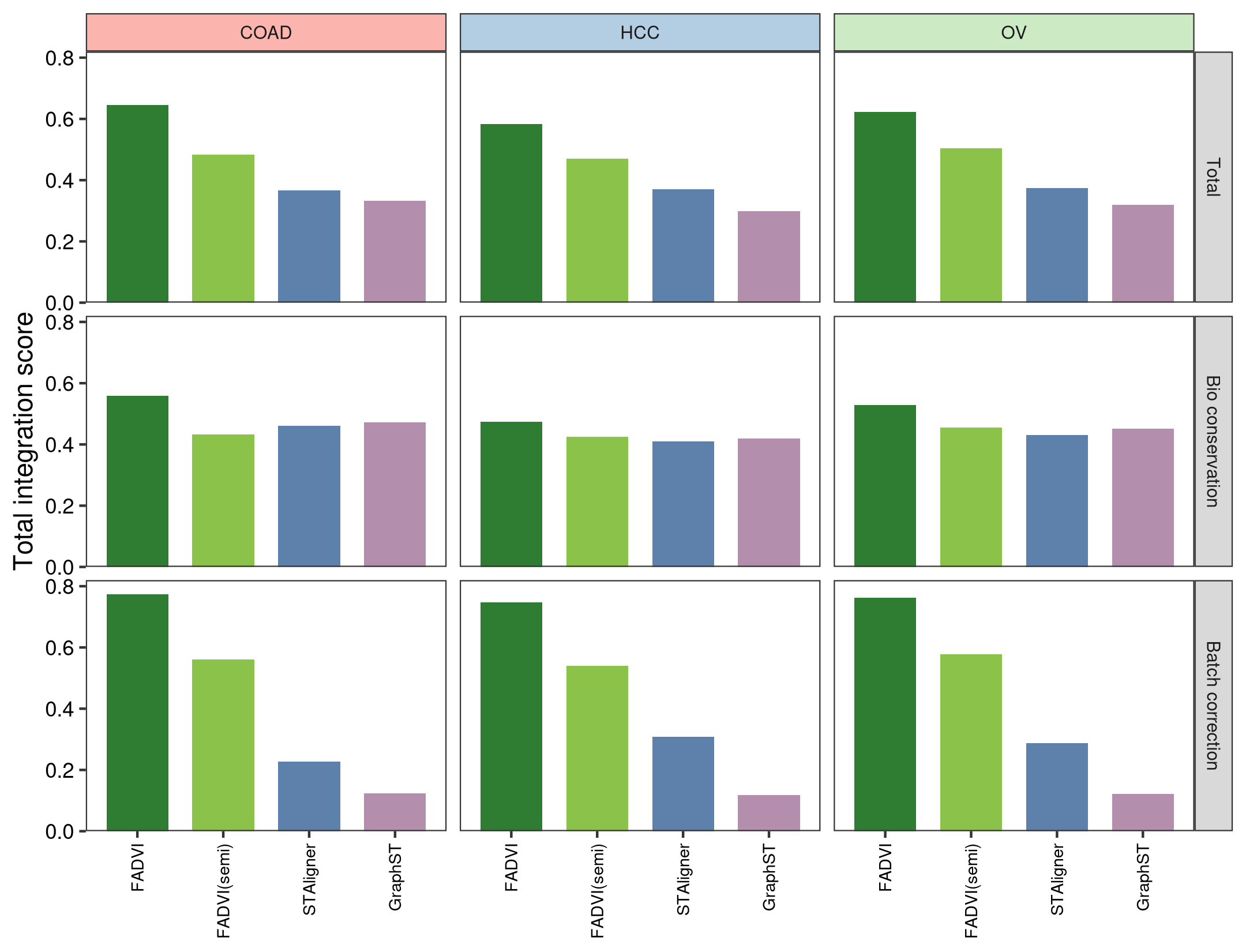

**Figure S22. FADVI outperforms spatially-aware integration methods that explicitly use spatial coordinates for cross-platform integration.** Total, bio conservation and batch correction scores for FADVI, semi-supervised FADVI, STAligner and GraphST in the three ST datasets. GraphST constructs its neighbor graph from spatial coordinates and STAligner computes a per-slice spatial network. Because neither method could process the full datasets within available memory, all methods were compared on the contiguous center square crops of approximately 12,500 cells per platform, which preserves the local adjacency these methods depend on; the scRNA-seq reference has no coordinates and is excluded. FADVI achieves the highest total score in all three datasets, driven primarily by batch correction, while bio conservation is comparable across all four methods.
